# Quantitative Proteomic Profiling of Murine Embryonic Heart Development Reveals a Role for the Mevalonate Pathway in Cardiomyocyte Proliferation

**DOI:** 10.1101/2022.02.21.481309

**Authors:** Whitney Edwards, Todd M. Greco, Gregory E. Miner, Natalie K. Barker, Laura Herring, Sarah Cohen, Ileana M. Cristea, Frank L. Conlon

**Affiliations:** Department of Biology and Genetics, McAllister Heart Institute, UNC-Chapel Hill, Chapel Hill, NC 27599, USA; Department of Pharmacology, UNC-Chapel Hill, Chapel Hill, NC 27599, USA; Department of Molecular Biology, Princeton University, Washington Road, Princeton, NJ 08544, USA; Department of Cell Biology and Physiology, UNC-Chapel Hill, Chapel Hill, NC 27599, USA; Lineberger Comprehensive Cancer Center, University of North Carolina at Chapel Hill, Chapel Hill, NC 27599, USA

**Keywords:** cardiac, heart, proteomics, mevalonate pathway, metabolism, congenital heart disease, heart development

## Abstract

Defining the molecular mechanisms that govern heart development is essential for identifying the etiology of congenital heart disease. Here, quantitative proteomics was used to measure temporal changes in the cardiac proteome at eight critical stages of murine embryonic heart development. Global temporal profiles of the over 7,300 identified proteins uncovered signature cardiac protein interaction networks that linked protein dynamics with molecular pathways. Using this integrated dataset, we identified and established a functional role for the mevalonate pathway in the regulation of embryonic cardiomyocyte proliferation and cell signaling. Overall, our proteomic datasets are an invaluable resource for studying molecular events that regulate embryonic heart development and contribute to congenital heart disease.

## INTRODUCTION

Congenital heart disease (CHD) is the most common type of birth defect, affecting approximately 1% of live births (Gilboa et al., 2016; Hoffman and Kaplan, 2002; Reller et al., 2008). The structural malformations that characterize CHD result from perturbations in the molecular programs that regulate heart development; understanding the details of these mechanisms and how they go awry will be critical for better understanding cardiac development and addressing CHDs.

The susceptibility of the heart to congenital defects reflects the highly complex developmental processes required for proper heart formation, which include precise coordination of cardiac cell specification and differentiation with morphogenic events that guide development of the different heart structures. These processes must occur over a surprisingly short period. The heart begins as a linear heart tube, which subsequently undergoes cardiac looping, a key morphogenic event that is essential for proper orientation of the heart chambers. This developmental milestone is followed by maturation of the cardiac chambers, atrial and interventricular septation, development of the heart valves, and formation of the cardiac conduction system (Bruneau, 2002; Kelly et al., 2014; Kirby and Waldo, 2002; Savolainen et al., 2009). This transition from a linear heart tube to a 4-chambered heart that anatomically parallels the adult heart occurs within three weeks in humans and one week in mice (Krishnan et al., 2014).

Identifying the molecular mechanisms that regulate mammalian heart development is essential for understanding the etiology of CHDs. Traditionally, genomics and transcriptomic approaches have been used to identify key genes and regulatory networks that govern critical steps of cardiac development. Recent advances in single-cell RNA sequencing have enabled detailed analyses of cell-specific transcriptional profiles and lineage trajectories in cardiac development and disease (Paik et al., 2020; Samad and Wu, 2021). These methodologies have enhanced our knowledge of transcriptional dynamics; however, these dynamics likely do not accurately reflect protein expression dynamics in the developing heart. RNA transcript levels often correlate poorly with protein abundance because multiple post-translational mechanisms influence protein expression (Chick et al., 2016; Nie et al., 2007; De Sousa Abreu et al., 2009). Therefore, a comprehensive analysis of temporal protein expression dynamics is essential to fully understand the molecular mechanisms of heart development and CHD.

In this study, we used multiplexed quantitative proteomics to quantify protein abundances and assemble proteome profiles across eight time points of murine embryonic heart development. We quantified the temporal expression of 7,313 cardiac proteins, of which 3,799 proteins exhibited differential expression during development. Using these embryonic temporal proteome profiles of cardiac development, we linked protein dynamics with molecular pathways. In addition, we monitored the temporal protein expression of 185 congenital heart disease-associated proteins. Further, we identified and functionally investigate protein pathways that regulate embryonic heart development. Our analysis uncovered an unexpected overrepresentation of core components of the mevalonate (MVA) pathway during mid-gestation of heart development. Concordantly, our functional analysis demonstrated that the MVA pathway regulates embryonic cardiomyocyte proliferation and cell signaling. To our knowledge, we have generated the largest proteomic dataset for the murine embryonic heart, representing a critical resource for measuring protein dynamics, protein interaction networks, and regulatory pathways that are essential for embryonic heart development.

## RESULTS

### Quantification of temporal protein abundance during embryonic heart development

Identifying embryonic cardiac proteins and delineating temporal regulation of protein networks and pathways is critical for establishing molecular mechanisms that govern heart development. Here, we used quantitative mass spectrometry with Tandem Mass Tag labeling to investigate protein abundance dynamics at critical stages of heart development. This quantitative multiplexing approach is ideal for studying the embryonic heart proteome because it enables high-throughput quantification of proteins from relatively low amounts of tissues, while minimizing variability and missing values between samples and supporting detection of low abundance proteins (Pappireddi et al., 2019; Plubell et al., 2017; Zecha et al., 2019).

We collected hearts at eight stages, from embryonic day 9.5 (E9.5) to day 16.5 (E16.5) (n=3 biological replicates). These stages include essential cardiac morphogenetic events, namely, cardiac looping, chamber formation, atrial and ventricular septation, and valve development (Figure 1A). To obtain high-confidence candidates, we used stringent filtering parameters that resulted in the quantification of 7,313 proteins (Figure 1B). Pearson correlation between the three biological replicates averaged 0.98; thus, protein identification and quantification were highly reproducible (Figure S2). To identify differentially expressed proteins during development, protein abundance ratios were calculated for each protein relative to the E9.5 timepoint. A total of 3,799 proteins displayed significant temporal changes in abundance at one or more of the developmental stages (|Log_2_ fold change| ≥ 0.585 and adjusted p-value ≤ 0.05; Figures 1B, S3).

**Figure 1.**
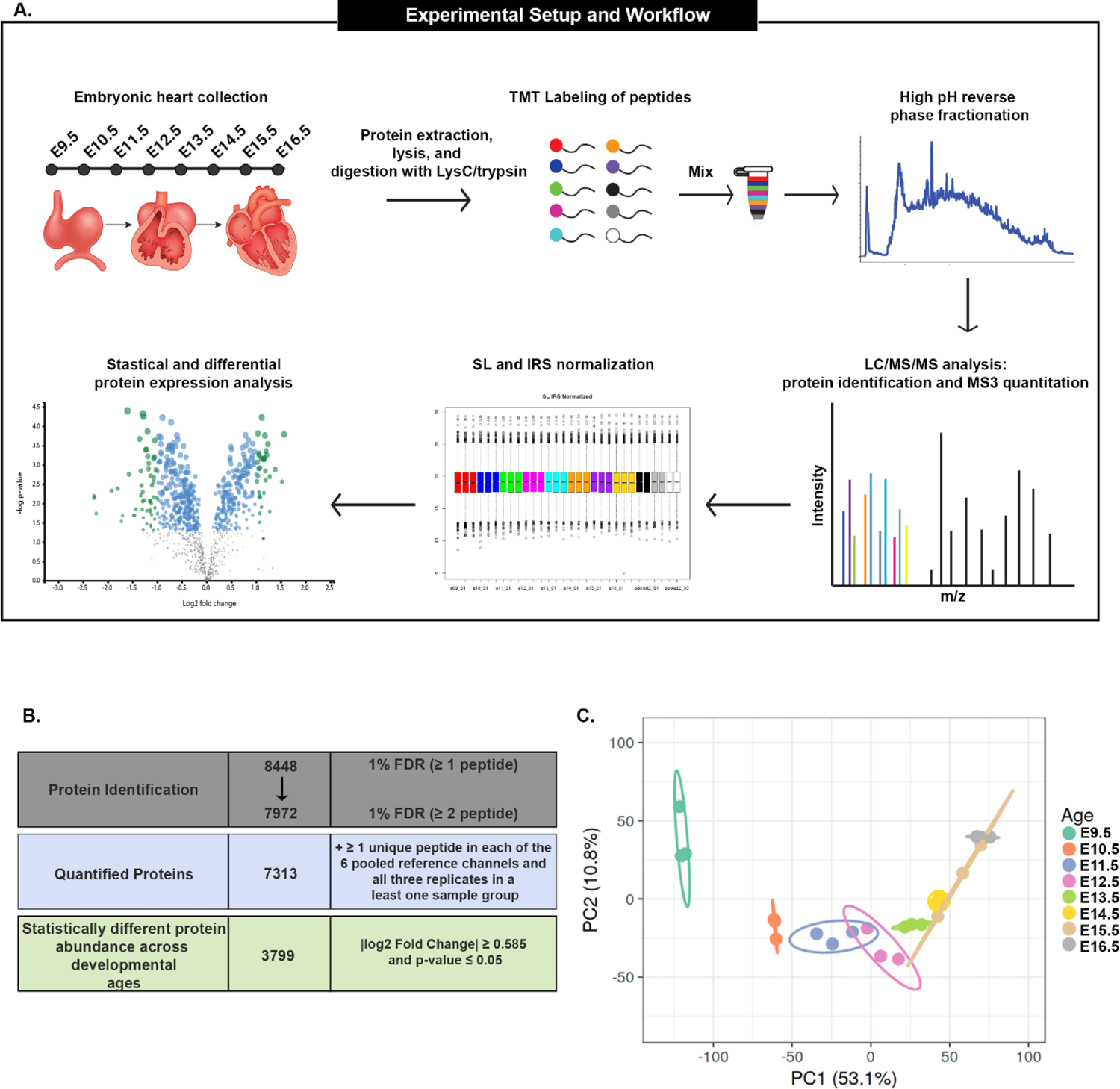
Quantification of temporal protein expression during embryonic heart development. (A) Tandem mass tag (TMT) quantitative mass spectrometry (MS) experimental design and analysis used to profile protein abundance in the developing mouse heart. Three independent TMT-MS experiments were performed. (B) Filtering parameters used to generate final list of quantified proteins (7,313) and statistical cutoffs used to determine differentially expressed proteins (3,799). (C) Principal component analysis (PCA) of TMT-MS proteomic data from each of the developmental time points. Color indicates age. Ellipses represent 95% confidence intervals.

Principal component analysis (PCA) showed separation of the embryonic stages, while the biological replicates of the same age clustered closely together (Figure 1C). This PCA agrees with known differences in morphological/developmental events between these stages of embryonic heart development. For example, the E9.5 timepoint differed the most from the other ages, because, at this age, the heart has yet to undergo key developmental processes such as cardiac chamber maturation and valve formation. In addition, the E15.5 replicates clustered closely with the E14.5 and E16.5 samples, which suggested a strong similarity in protein expression at these ages when cardiac morphogenesis is nearly complete. To our knowledge, this dataset represents the largest comprehensive analysis of proteins abundance dynamics during embryonic heart development.

### Expression dynamics of cell-type-specific proteins in the developing heart

Heart development and function rely on the coordinated maturation of many cardiac cell types, such as cardiomyocytes, endocardial cells, and valve endothelial cells. We performed a targeted analysis to assess whether our proteomic dataset identified proteins associated with specific cardiac cell types and whether the data were consistent with previously demonstrated temporal protein dynamics.

First, we examined proteins associated with cardiomyocyte (CM) development and function. During embryogenesis, the rapid increase in CM number is accompanied by the initiation of CM maturation. Cardiomyocyte maturation coincides with significant structural changes and extensive reorganization of the cardiac contractile machinery (de Boer et al., 2012; Guo and Pu, 2020; Hirschy et al., 2006). Concordantly, we observed a significant and steady increase in CM-specific proteins and cardiac contractile proteins between E9.5 and E16.5. These proteins included troponin complex proteins (TMP1, TNNT2), myofibril proteins (MYH7 and MYL2) and well-established cardiomyocyte-specific markers (NPPA, SMPX) (Figure 2A).

**Figure 2.**
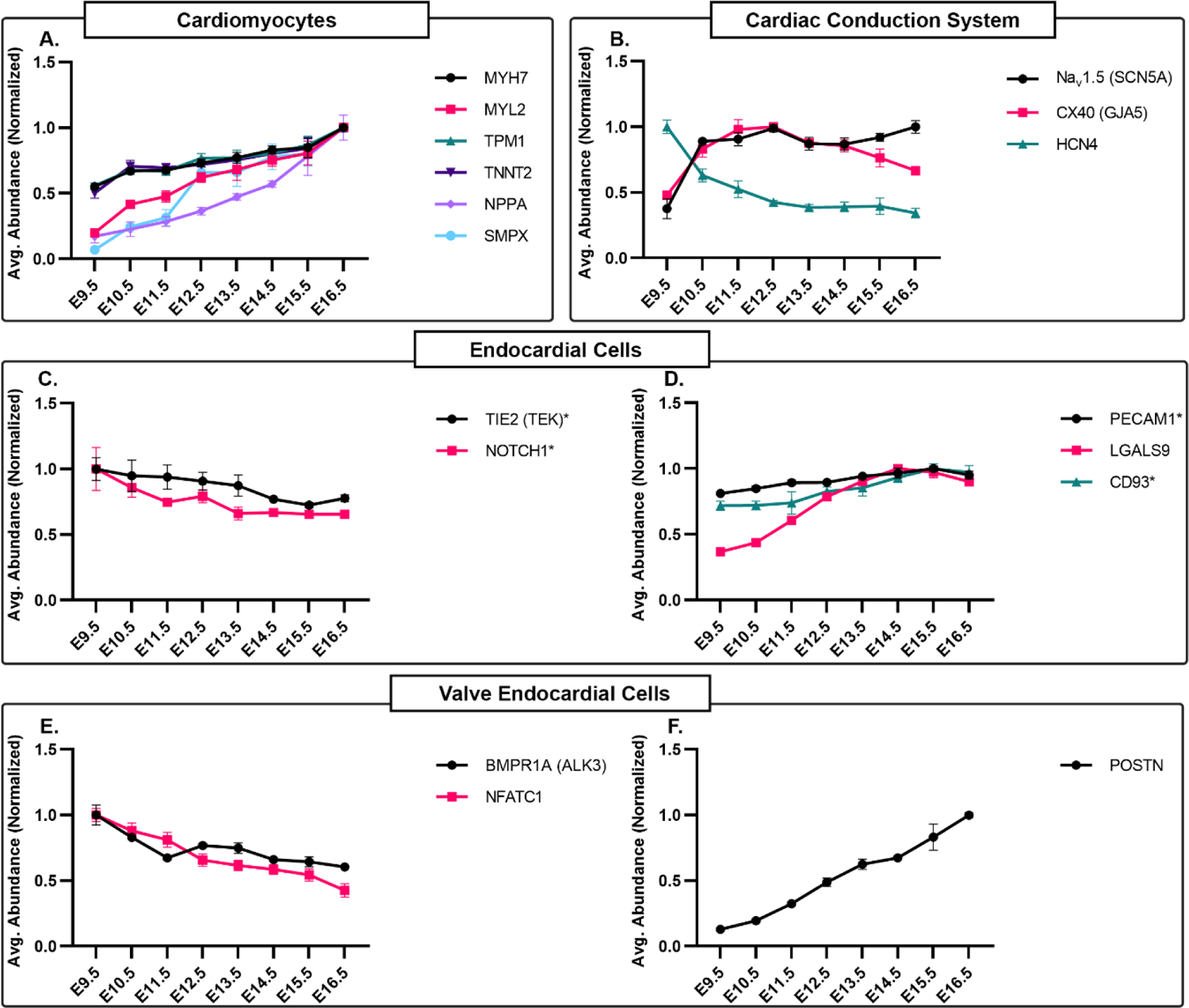
Expression dynamics of cell type-specific proteins in the developing heart. Temporal expression profiles of proteins specific to cardiomyocytes (A), cardiac conduction system (B), endocardial cells (C, D), and valve endocardial cells (E, F). For panels A-F, graphs represent average protein abundance (maximum average normalized) at each development point. Error bars represent SEM. Asterisk (*) indicates proteins that met statistical but not fold change cutoffs for differential expression.

We also profiled proteins associated with maturation and function of the cardiac conduction system (CCS), a specialized group of CMs that generate and propagate electrical impulses through the heart. The CCS is a complex system of functionally distinct structures including the sinoatrial node (SAN), atrioventricular node, atrioventricular bundle, bundle branches, and the Purkinje fibers. Critical steps of CCS cell differentiation and maturation occur between E9.5 and E16.5. Therefore, we used our dataset to identify and characterize the expression dynamics of well-established CCS-associated proteins. The ion channel HCN4 is required for cardiac pacemaker activity of the SAN; HCN4 exhibits broad expression in the myocardium early in development and becomes progressively more restricted to CCS cells by the end of embryogenesis (Liang et al., 2013, 2015; Wu et al., 2014). This expression pattern was mirrored by a gradual decrease in HCN4 expression from E9.5 to E16.5 (Figure 2B). In contrast, CX40 (GJA5), a gap junction protein associated with the ventricular portion of the mature CCS, showed a drastic increase in expression from E9.5 to E10.5, maintaining a relatively steady expression level during mid-gestation (E11.5-E13.5), followed by a decrease in expression at later stages (Figure 2B). This expression pattern of CX40 agreed with immunohistological studies of CX40 in the embryonic heart (Coppen et al., 2003; Delorme et al., 1995).

We next examined proteins associated with endocardial cells (EC) which are specialized endothelial cells that form the innermost layer of the heart. These cells contribute to critical processes during heart development including promoting the formation of the valves and maturation of the trabecular layer of the myocardium. First, we analyzed protein expression dynamics of broad EC cell markers. Previous studies demonstrated that TIE2 (TEK) and NOTCH1 are expressed throughout the endocardium during development, and the absence of either marker causes attenuated endocardial cell proliferation and impaired trabeculation, ultimately resulting in embryonic lethality by E10.5 (Dumont et al., 1994; Grego-Bessa et al., 2007; Qu et al., 2019; Sato et al., 1995; Timmerman et al., 2004). Our analysis showed that NOTCH1 and TIE2 displayed relatively steady expression levels with only minimal decreases observed at later gestational stages (E14.5-E16.5) (Figure 2C). PECAM1 (CD31), another well-established broad EC marker, also showed minimal changes in expression level between E9.5 and E16.5. We also examined the expression of two recently identified EC-specific proteins, LGALS9 and CD93. CD93 parallels the expression pattern of PECAM1, while LGALS9 shows a substantial increase in expression between E9.5 and E13.5. (DeLaughter et al., 2016) (Figure 2D).

Valve endothelial cells (VEC) are a subset of EC cells that are critical for embryonic valve development. At E9.5, VECs contribute to the formation of the cardiac cushions (primordial valves). Between E9.5 and E16.5, the cardiac cushions undergo extensive growth and remodeling as the valves mature (Tao et al., 2012). We first examined VEC markers that are required for the initial steps of cardiac cushion formation. NFATC1, a transcription factor enriched in VECs, is associated with congenital malformations of the cardiac valves in mice and humans (Abdul-Sater et al., 2012; Ranger et al., 1998; Wu et al., 2011, 2013). We observed that NFATC1 displayed peak protein abundance at E9.5 and steadily decreased as heart development proceeded (Figure 2E). We also examined BMPR1A, a BMP receptor that has a well-established function in epithelial-to-mesenchymal (EMT) transition in the primordial valve (Gaussin et al., 2005; Ma et al., 2005). Our data showed that BMPR1A had the highest abundance at E9.5 and significantly decreased at later stages of embryogenesis (Figure 2E). In contrast, POSTN, an extracellular matrix protein required for valve maturation, showed a marked increase in abundance between E9.5 and E16.5 (Figure 2F). These data agreed with immunohistological studies that showed POSTN expression enriched in the cardiac valves as they mature (Norris et al., 2008).

Collectively, we demonstrated that our dataset can be used to analyze time-resolved expression dynamics of cell-type-specific proteins in the developing heart. Furthermore, the accuracy and depth of our proteomic dataset are exemplified by its precise recapitulation of known protein expression patterns associated with critical developmental processes in the heart.

### Chromatin organization and energy metabolism cardiac protein networks are modulated from early-to-mid gestation

To determine the molecular pathways and protein networks that are differentially regulated during cardiac development, we performed pairwise comparisons of the cardiac proteomes of distinct embryonic stages. Progression of heart development from E9.5 to E11.5 represents the beginning of many key developmental events, including chamber differentiation/growth and the onset of valve formation and chamber septation. To define the protein networks involved in these processes, we compared differences in protein expression between E9.5 and E11.5. We identified 1,559 differentially expressed proteins, of which 515 proteins showed significantly higher abundance at E9.5, and 1,044 proteins displayed higher abundance at E11.5 (|Log_2_ fold change| ≥ 0.585 and adjusted p-value ≤ 0.05; Figure 3A).

**Figure 3.**
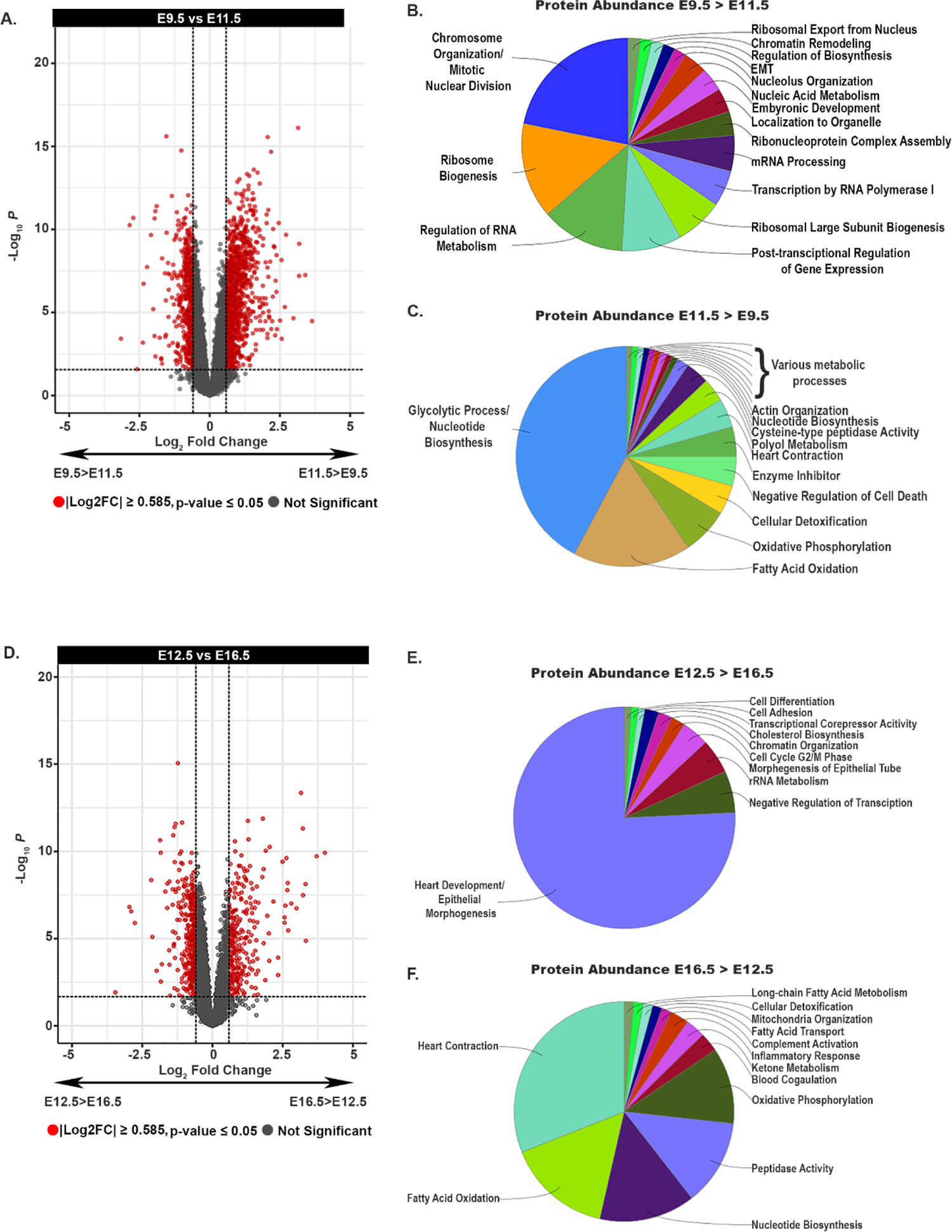
Comparison of cardiac proteome at distinct embryonic stages. (A) Volcano plot displaying protein abundance differences between E9.5 and E11.5. (B) Representation of significantly enriched (p-value ≤ 0.001) GO terms (biological process) associated proteins that display greater abundance at E9.5 compared with E11.5. (C) Representation of significantly enriched (p-value ≤ 0.001) GO terms (biological process) associated proteins that display greater abundance at E11.5 compared with E9.5. (D) Volcano plot displaying protein abundance differences between E12.5 and E16.5. (E) Representation of significantly enriched (p-value ≤ 0.001) GO terms (biological process) associated proteins that display greater abundance at E12.5 compared with E16.5. (F) Representation of significantly enriched (p-value ≤ 0.001) GO terms (biological process) associated proteins that display greater abundance at E16.5 compared with E12.5. For panels A and D: Plots are shown as Log2 fold change (x-axis) plotted against the -log10 of the p-value (y-axis). Red dots represent proteins that were considered significantly different based on fold change and p-value parameters.

Gene ontology (GO) pathway analysis revealed that proteins highly expressed at E9.5 were primarily associated with chromosome organization/mitotic nuclear division, ribosome biogenesis, and RNA transcription/processing (Figure 3B). We conducted STRING analyses to visualize functional protein interaction networks and to identify physical protein-protein interactions within select pathways. Proteins involved in chromatin organization formed a network that contained components of chromatin-modifying complexes including, Polycomb repressive complex I (PCGF6, RNF2, L3MBLT2, EHMT1), BAF complex (SMARCB1, SMARCC1, SMARCD1), and MOZ/MORF complex (KAT6A, ING5) (Figure S4A). These multi-subunit complexes are epigenetic modifiers that regulate cardiac gene expression programs.

Genetic deletion of specific components in each of these complexes is associated with severe anomalies in cardiac morphogenesis (Hota and Bruneau, 2016; Hota et al., 2019; Lickert et al., 2004; Shirai et al., 2002, 2016; Sun et al., 2018; Takihara et al., 1997; Vanyai et al., 2015; Voss et al., 2012). For example, loss of BAF complex subunits results in impaired cardiomyocyte differentiation and embryonic lethality by E10.0 (Lickert et al., 2004). Additionally, mutations in components of the BAF, PRC1, and MOZ/MORF complexes were recently identified in a study that classified de novo damaging mutations in isolated CHD cases; these findings illustrated a critical function for these complexes in human heart development (Jin et al., 2017).

Our analyses identified a second large interaction network of proteins associated with cell cycle control. This set of proteins included Kinesin family members (KIF20A, KIFB20B, KIF23) and proteins that regulate mitotic spindle formation (AURKB, PLK1, SPDL1, INCENP) (Figure S4B). These findings were consistent with E9.5 hearts undergoing a period of rapid growth and division.

Interestingly, we also observed a large network of ribosome biogenesis proteins upregulated at E9.5. This network included ribosomal subunits (RPS19, RPL11, RPL35A) and proteins involved in ribosomal RNA (rRNA) processing (WDR43, WDR75, UTP18) (Figure S4C). Although the particular function for ribosome biogenesis in heart development is not well understood, mutations in proteins involved in ribosome biogenesis are associated with congenital heart defects and developmental disorders referred to as ribosomopathies (Farley-Barnes et al., 2019; Venturi and Montanaro, 2020). The most well-studied ribosomopathy is Diamond-Blackfan anemia, a congenital syndrome caused by mutations in ribosomal subunit proteins that presents with a wide range of heart defects including ventricular and atrial septal defects and Tetralogy of Fallot (Vlachos et al., 2018).

In contrast to our findings at E9.5, the cardiac proteins highly expressed at E11.5 were associated with metabolic pathways including energy metabolism, fatty acid oxidation, and glycolysis (Figure 3C). Enriched proteins at E11.5 formed highly interconnected protein-protein interaction networks composed of proteins associated with nucleotide biosynthesis (AK1, NT5C, PDE4D), glycolysis (ALDOA, GAPDH, PGK1), fatty acid oxidation (ACAA2, ETFA, HADHA), and lipid peroxidation (GPX1, PRDX6, GSTM7) (Figure S5B and S5C). In addition, we observed an increase in complexes associated with heart contraction, including the dystrophin-associated complex (DMD, SGCD, CAV3) and sarcomeric proteins (TNNI3, MYL2, TCAP) (Figure S5A). The increase in cardiac protein networks associated with metabolic pathways and cardiac muscle function indicates the maturation and growth of the myocardium, which requires increased energy metabolites at these stages.

### TGFB/WNT signaling, transcriptional repression and cardiac muscle protein networks are modulated from mid-to-late gestation

Heart development between E12.5 to E16.5 is the final phase of critical morphogenic events. During this developmental window, the heart chambers are septated, the primordial valves undergo extensive remodeling, and by E16.5 the heart morphologically reflects the postnatal heart. Our analyses of E12.5 versus E16.5 revealed enriched biological processes distinct from those identified in the E9.5 versus E11.5 comparison. Overall, we identified 679 differentially expressed cardiac proteins, of which 397 proteins had significantly higher abundance at E12.5, and 283 proteins displayed higher abundance at E16.5 (Figure 3D).

Biological pathways enriched at E12.5 included epithelial morphogenesis, negative regulation of transcription, and rRNA metabolism (Figure 3E). Proteins associated within the epithelial morphogenesis category formed functional protein interaction networks that included members of the transforming growth factor beta (TGFB) (TGFB1, TGFBR1, TGFBR2,) and WNT (WNT5A, FZD1, FZD7) signaling pathways (Figure S6A and S6B). The TGFB and WNT signaling pathways have multifaceted functions in embryonic heart development. Therefore, enriched expression of proteins associated with these pathways at E12.5 may reflect their functions in valve maturation, trabeculation, and/or septation of the vessels and chambers. We also identified a large interaction network associated with transcriptional repression.

Interestingly, similar to E9.5, we identified a subset of cardiac proteins associated with the Polycomb Repressive Complexes (SUZ12, EZH2, RYBP; Figure S6C). Notably, many of these proteins were distinct from those identified at E9.5, which suggested systematic temporal regulation of distinct Polycomb Repressive complex proteins during cardiac development (Figure S4A). In comparison, enriched pathways and protein interaction networks at E16.5 were associated predominantly with cardiac muscle function and included proteins involved in heart contraction (DMD, TNNI3, MYL2), fatty acid oxidation (ACAA2, HADHB, EFTA), and oxidative phosphorylation (COX5A, NDUFA4, UQCRH) (Figure 3F, S7A-C).

In sum, these quantitative proteomic analyses identified key proteins, biological pathways, and protein complexes that are differentially regulated during essential phases of mammalian heart development. Additionally, this work illustrates how our dataset can be used to conduct targeted analyses of age-specific changes in the embryonic cardiac proteome.

### Analysis of global protein abundance reveals eight distinct temporal cardiac protein expression patterns

A unique advantage of our dataset is that it includes the cardiac proteome profiles of eight embryonic time points which encompass a wide range of cardiac development milestones. Therefore, we sought to map global changes in protein expression across the mammalian cardiac developmental timescale. Hierarchical clustering was performed on 3,799 proteins that were differentially expressed from E9.5 to E16.5 (Figure 1B). We uncovered eight distinct cardiac protein expression profiles (Clusters; Figure 4A).

**Figure 4:**
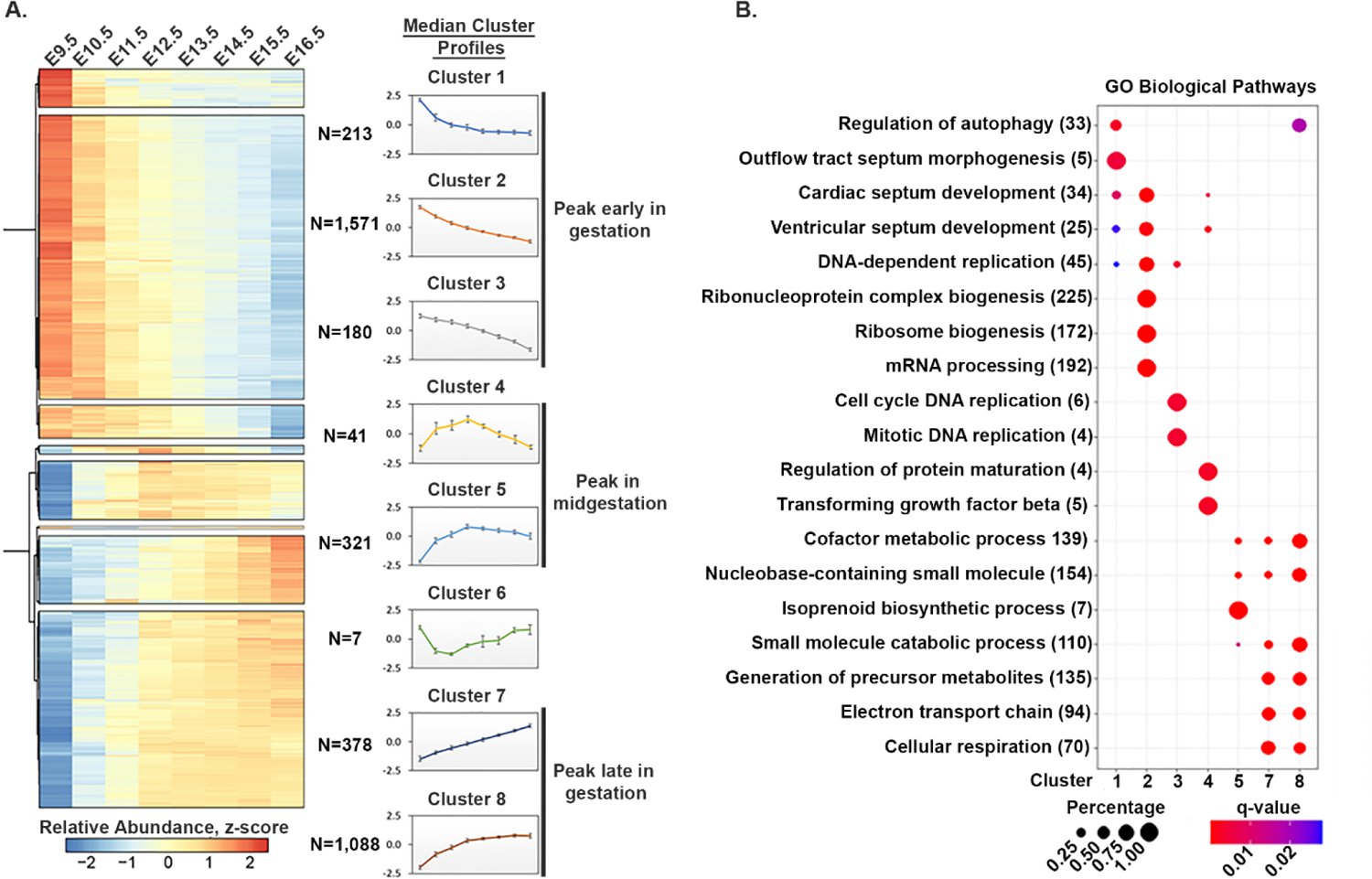
Analysis of global protein abundance reveals eight distinct temporal cardiac protein expression patterns. (A) Hierarchical clustering of 3,799 differentially expressed proteins revealed eight distinct temporal patterns of protein expression (clusters 1-8). For each developmental timepoint the median values of all the proteins within a cluster were calculated to generate the median cluster profile graphs. (B) Dot plot of GO terms (biological processes) significantly enriched for each cluster. Color indicates q-value. Size of circles indicates percent of input proteins in a given biological process term.

Clusters 1, 2, and 3 displayed peak protein abundance early in gestation (E9.5-E10.5). Cluster 1 proteins showed a rapid decrease at E11.5 and maintained steady expression at later stages in development. Proteins in Cluster 1 (N=213) were enriched for pathways involved in autophagy, outflow tract morphogenesis, and septum development (Figure 4B). In contrast, Clusters 2 and 3 showed a steady decrease in abundance as development progressed. We note that Cluster 2 contained the largest number of proteins (N=1571), accounting for approximately 41% of all differentially expressed proteins. Pathways specifically associated with Cluster 2 included ribosome biogenesis and mRNA processing, (Figure 3B). Cluster 3 proteins (N=180) were linked to pathways associated with DNA replication and cell cycle progression. These results paralleled pathways we identified in the E9.5 versus E11.5 pairwise comparison (Figure 3A-C).

Clusters 4 (N=41) and 5 (N=321) included proteins that exhibited the highest expression during mid-gestation (E11.5-E13.5). Cluster 4 proteins displayed a sharp increase in expression between E9.5 and E12.5 followed by an immediate and continuous decrease at later stages. This group was enriched for pathways involved in protein maturation and TGFB signaling. Cluster 5 proteins showed a similar pattern of expression and were specifically associated with metabolic pathways, including cofactor metabolic processes, small molecule metabolic processes, and isoprenoid biosynthesis.

In contrast to Cluster 4 and 5, proteins in Cluster 6 displayed the opposite expression pattern, with high abundance at E9.5, followed by a sharp and transient decrease in expression by E11.5 and then a gradual increase during late gestation. Because of the small number of proteins in Cluster 6 (N=7), GO pathway analysis was unable to statistically correlate this group with any specific biological processes, however, we noted that several proteins were associated with the extracellular matrix (ITGA7, COL5A1, COL1A2).

Proteins in clusters 7 (N= 378) and 8 (N=1088) showed peak protein abundance in late gestation (E13.5-E16.5; Figure 4B). Proteins in Cluster 7 and 8 were enriched for pathways associated with metabolism, the electron transport chain, and respiration.

Overall, our dataset enabled us to analyze global changes in cardiac protein expression during embryonic heart development. Our identification of eight distinct protein expression profiles suggests that strict temporal regulation of protein abundance is essential for embryonic heart development. Further investigation of individual proteins and protein networks within each of these temporally regulated pathways will provide new molecular explanations of cardiac development.

### Expression dynamics of congenital heart disease-associated proteins

Clinical and genetic studies have identified a large cohort of genes that are implicated in congenital heart disease (CHD). We used our datasets to investigate the temporal expression of cardiac proteins associated with isolated and syndromic CHDs. To conduct these analyses, we curated a list of 185 proteins commonly implicated in CHDs in mice and humans (Fahed et al., 2013; Jin et al., 2017; Nees and Chung, 2020; Williams et al., 2019; Zaidi and Brueckner, 2017). Most (100/185) of the CHD-associated proteins were differentially expressed during the course of embryonic heart development (Figure 5). Strikingly, approximately 71% of the differentially expressed proteins displayed peak abundances at early development time point (Clusters 1, 2, 3). These proteins were associated with signaling pathways (JAG1, SMAD4, TGFBR1), chromatin modification (CHD4, CHD7, JARID2), cardiac transcription factors (TBX5, GATA4, NKX2-5), and ribosomal proteins (RPS19, RPS17, RPL11). A smaller subset of proteins displayed peak abundance during mid (11/100; Cluster 4, 5) and late gestation (16/100; Cluster 7, 8). This smaller group included several structural (ELN, FBN1, MYH7) and intracellular signaling proteins (MAP2K1, PTEN, SOS2). Interestingly, included in Cluster 6 were two collagen proteins associated with CHDs (COL5A1 and COL1A2).

**Figure 5:**
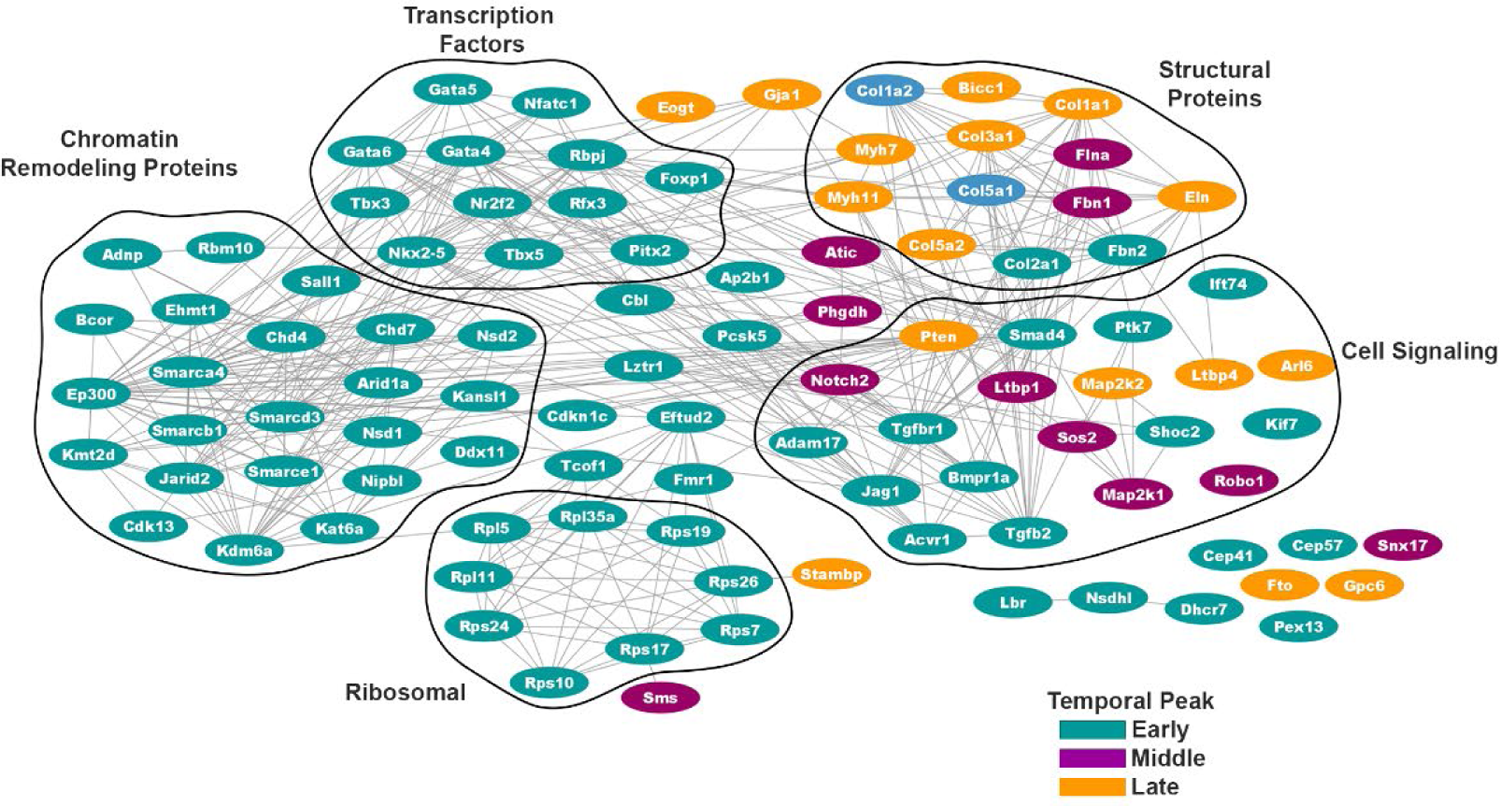
Expression dynamics of CHD-associated proteins. Functional protein interaction network of 100 differentially expressed CHD-associated proteins grouped by functional categories. Interaction network is based on function and physical protein interactions. Teal nodes= proteins with peak abundance in early gestation (cluster 1, 2, 3), purple nodes= proteins with peak abundance in mid-gestation (cluster 4, 5), orange nodes= proteins with peak abundance in late gestation (cluster 7, 8), Light blue nodes= proteins with a decrease in protein abundance during mid-gestation (cluster 6).

Overall, our dataset provided temporal protein dynamics for a large cohort of CHD-associated proteins. Furthermore, our analysis demonstrates that the majority these proteins display peak protein abundance at early stages of heart development (E9.5-E10.5) indicating that this embryonic window is particularly sensitive to disruption in protein abundances.

### Mevalonate pathway proteins are abundantly expressed in mid-gestation

One of our goals in generating an extensive proteomic dataset was to identify novel and less well-studied proteins and pathways that are essential for heart development. We were specifically interested in proteins that displayed non-monotonic expression patterns. Thus, we further investigated proteins represented in Clusters 4 and 5 that showed a peak in protein abundance at mid-gestation (Figure 4A, 4B). Intriguingly, most of the core enzymes of the mevalonate (MVA) pathway (aka, isoprenoid biosynthesis pathway) were found in Cluster 5. These enzymes included MVK, PMVK, MVD and FDPS (Figure 6A, 6B). The MVA pathway is an essential metabolic pathway that converts acetyl-CoA to either cholesterol or lipid moieties for protein prenylation. This pathway regulates a diverse set of cellular functions, which include proliferation, apoptosis, and cell migration (Gong et al., 2019; Mullen et al., 2016). Although relatively little is known about the function of the MVA pathway in the embryonic heart, previous studies suggest this pathway regulates cardiomyocyte proliferation and cytoarchitectural integrity (Chen et al., 2018; Mills et al., 2019). In addition, several MVA pathway components and proteins prenylated by this pathway are associated with congenital and adult cardiovascular diseases in mice and humans (Table 1). For example, cardiac-specific deletion of Farnesyl Diphosphate Synthase (FDPS) in mice results in cardiac hypertrophy (Wang et al., 2021). In addition, multiple syndromes that present with congenital heart defects, such as Costello, Noonan, and Cardio-facio-cutaneous syndromes, result from mutations in RAS family member proteins that require prenylation for function (Calcagni et al., 2017; Jhang et al., 2016). These observations suggested that the MVA pathway is important for heart development and function.

**Figure 6:**
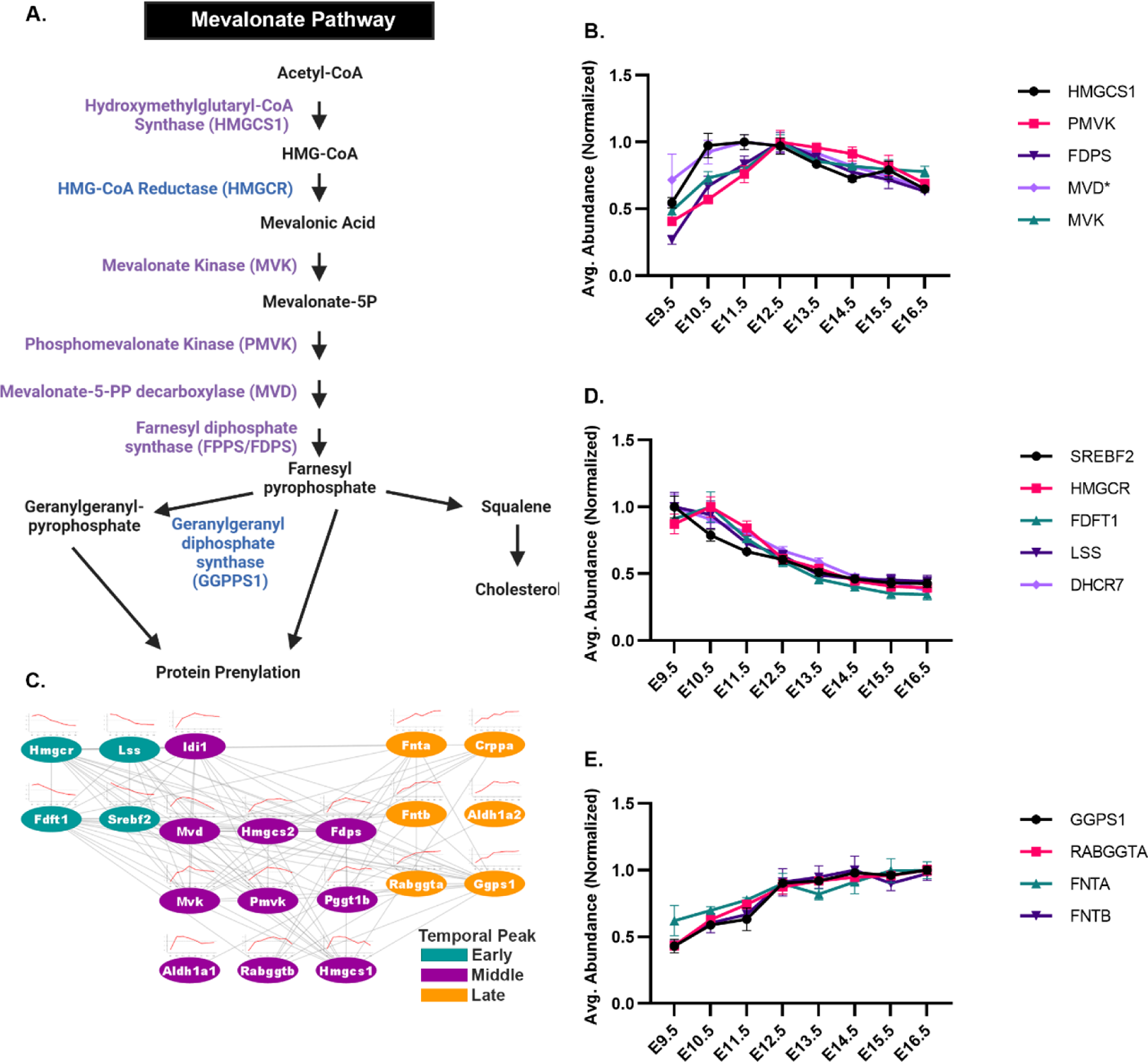
The mevalonate (MVA) pathway proteins are abundantly expressed in mid-gestation. (A) Schematic representation of the MVA pathway. Enzymes labeled in purple display peak abundance in mid-gestation (cluster 5). (B) Temporal protein expression profiles of MVA pathway-associated proteins that display peak abundance at mid-gestation (cluster 5). (C) STRING protein interaction network analysis performed on proteins associated with the MVA pathway. Interaction network is based on function and physical protein interactions. Teal nodes= proteins with peak abundance in early gestation (cluster 1, 2, 3), purple nodes= proteins with peak abundance in mid-gestation (cluster 4, 5), orange nodes= proteins with peak abundance in late gestation (cluster 7, 8). (D) Temporal protein expression profiles of MVA pathway-associated proteins that display peak abundance at early time point (cluster 2). (E) Temporal protein expression profiles of MVA pathway-associated proteins that display peak abundance at late time points (cluster 8). For panels B, D and E, graphs represent average protein abundance (maximum average normalized) at each development point. Error bars represent SEM. Asterisk (*) indicates proteins that met statistical but not fold change cutoffs for differential expression.

**Table 1:**
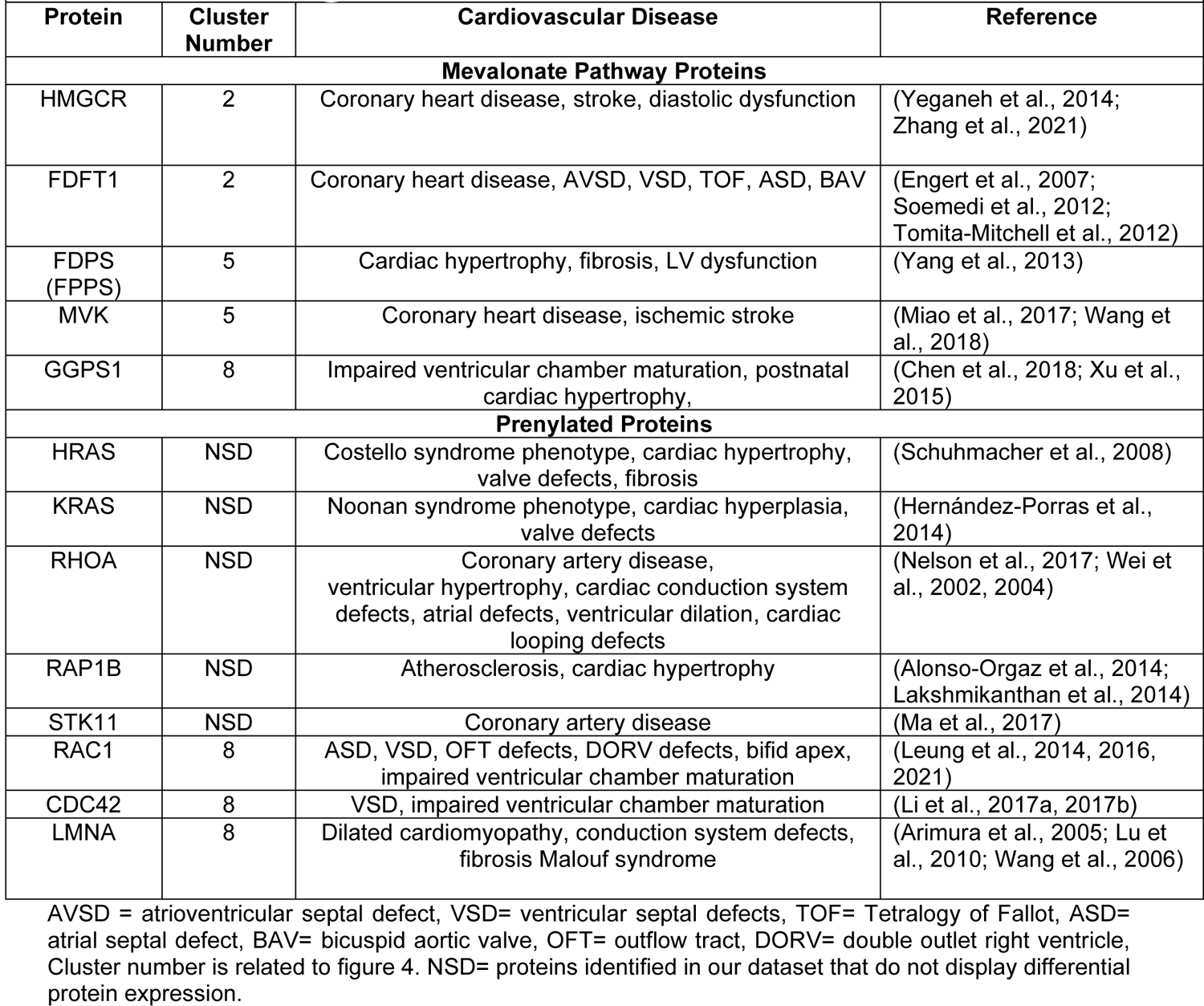
Mevalonate Pathway Proteins Associated with Cardiovascular Disease

We queried other Clusters to determine the expression patterns of additional MVA pathway components (Figure 6C). SREBF2 (SREBP2), a transcription factor that directly regulates the expression of MVA genes involved in cholesterol synthesis, was highly expressed at E9.5 and showed a steady decrease throughout development (Cluster 2). In addition, HMGCR1, FDFT1 (SQS), and LSS, transcriptional targets of SREBF2, all showed a similar decrease in expression (Figure 6D) (Sakakura et al., 2001). In contrast, pathway components that are specifically involved in the terminal steps of protein prenylation (GGPS1, RABGGTA, FTNA and FNTB) showed a marked increase in abundance, with peak abundance occurring at the end of gestation (Cluster 8) (Figure 6E). Of note, we observed opposing patterns of expression for MVA components that are directly involved in cholesterol biosynthesis (FDFT1, LSS, DHCR7) versus components that facilitate protein prenylation (GGPS1, FTNA, FNTB, RABGGTA). These results implied that different branches of the MVA pathway are active at distinct stages of embryonic heart development. In sum, our comprehensive dataset enabled us to identify and quantify temporal abundance profiles for key proteins in the cardiac MVA pathway.

### The mevalonate pathway controls cardiomyocyte proliferation and cell signaling pathways

Our temporal analysis of cardiac MVA pathway components showed that most of the core enzymes in the pathway displayed peak expression during mid-gestation (Cluster 5; Figure 4B, 6A, 6B). Therefore, we sought to define a function for the MVA pathway during mid-gestation of heart development. Previous studies suggest the MVA pathway regulates cardiomyocyte proliferation (Chen et al., 2018; Mills et al., 2019). However, no studies have tested direct control of embryonic cardiomyocyte proliferation by the MVA pathway.

We used a primary cardiomyocyte culture system to determine whether alterations in activity of the MVA pathway directly controlled cardiomyocyte proliferation. Statins are a large class of drugs that inhibit HMG-CoA reductase, the rate-limiting enzyme in the MVA pathway. Statins efficiently reduce cholesterol biosynthesis and production of isoprenoids used for protein prenylation. To measure effects of MVA pathway inhibition on embryonic cardiomyocyte proliferation, we isolated and treated E12.5 cardiomyocytes in culture for 24 hours with 10 µM simvastatin (SMV), a lipophilic statin shown to affect proliferation of human pluripotent stem cell-derived cardiomyocytes (Figure 7A) (Mills et al., 2019). Immunohistological analysis of cardiomyocyte cultures with the proliferation marker Ki67 revealed an almost 40% decrease in the number of Ki67 positive cardiomyocytes after SMV treatment compared with control (Figure 7B and 7C). No changes in overall cardiomyocyte number were detected with SMV treatment, which indicated that the decrease in proliferation was not due to cell death (Figure 7D). These results demonstrated that the MVA pathway can directly and acutely regulate embryonic cardiomyocyte proliferation.

**Figure 7:**
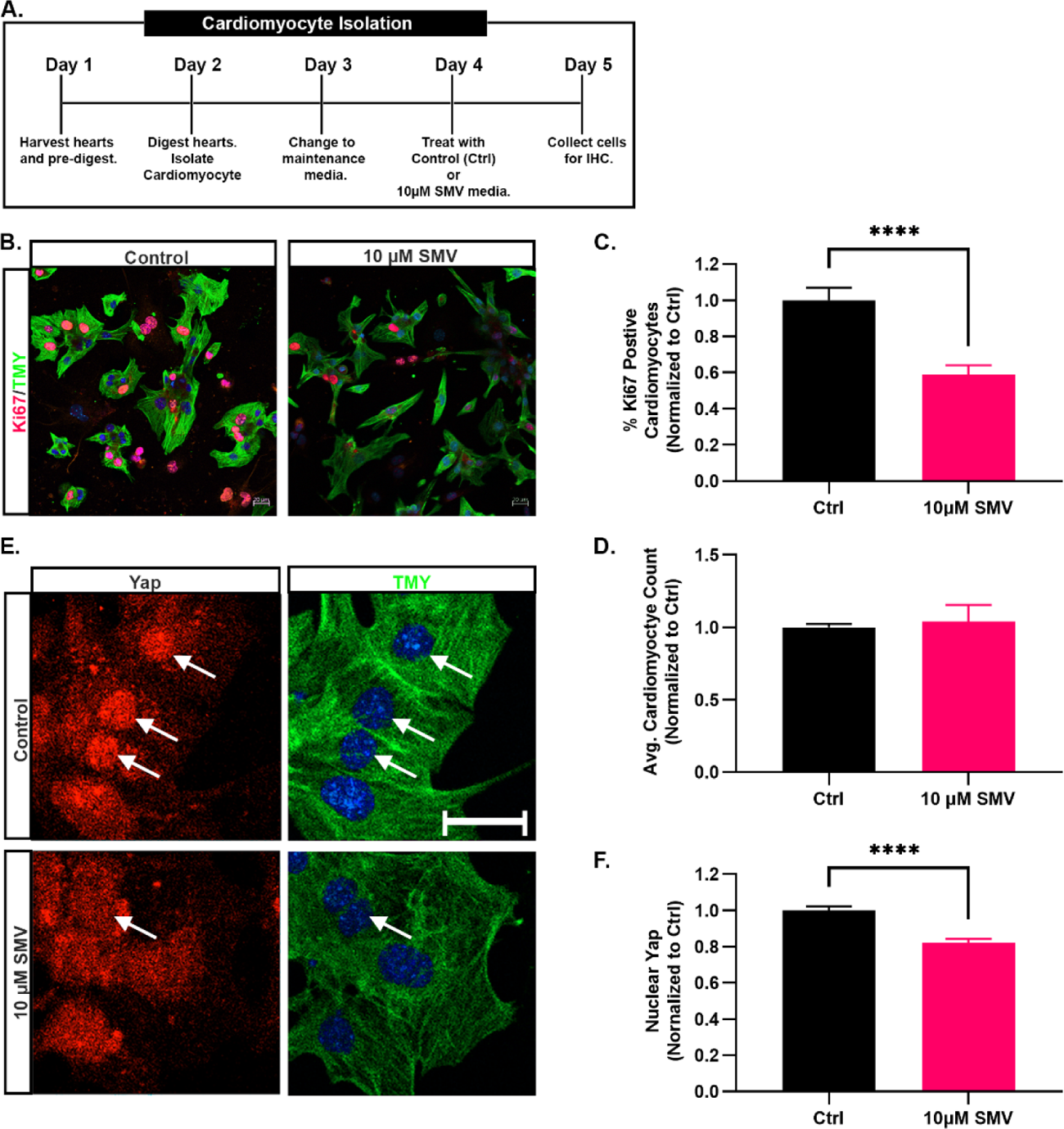
The mevalonate pathway controls cardiomyocyte proliferation and cell signaling pathways. (A) Schematic of embryonic cardiomyocyte isolation protocol and simvastatin (SMV) treatment paradigm. (B) Immunohistochemical analysis of proliferating (red; Ki67) cardiomyocytes (green; Tropomyosin) following treatment in culture with 10 µM SMV for 24 hours. TMY= tropomyosin. (C) Quantitation of percent of Ki67 positive cardiomyocytes following treatment in culture with 10 µM SMV for 24 hours. (D) Average number of cardiomyocytes following treatment in culture with 10 µM SMV for 24 hours. (E) Immunohistochemical analysis of nuclear YAP (red) accumulation in cardiomyocytes (green; Tropomyosin) following treatment in culture with 10 µM SMV for 24 hours. (F) Quantitation of average nuclear YAP intensity in cardiomyocytes following treatment in culture with 10 µM SMV for 24 hours. For panels C, D and F, graphs show values normalized to control. Scale bar=20 µm. Data are presented as ± SEM. **** denote p-value ≤ 0.0001 using a Student’s t-test.

YAP and TAZ are transcriptional mediators of the Hippo signaling pathway, a critical regulator of embryonic heart growth and maturation. Previous cancer studies showed that the MVA pathway promoted nuclear localization of YAP and TAZ, resulting in increased cell proliferation (Gong et al., 2019; Mullen et al., 2016; Sorrentino et al., 2014; Wang et al., 2014). Although YAP and TAZ have an established function in embryonic heart growth, regulation of these factors by the MVA pathway had not been shown in the embryonic heart (Chen et al., 2020; Moya and Halder, 2019). Therefore, we examined the effects of MVA pathway inhibition on YAP nuclear localization in embryonic cardiomyocytes. Using our primary cardiomyocyte culture paradigm, we found that E12.5 cardiomyocytes treated with SMV showed a significant reduction in nuclear YAP localization (Figure 7E and 7F). This finding demonstrated that the MVA pathway modulates YAP signaling in embryonic cardiomyocytes. Further, our results imply the MVA pathway controls cardiomyocyte proliferation by controlling YAP activity.

Collectively, our dataset enabled us to identify the temporal protein expression patterns of the MVA pathway components during embryonic heart development. Using this information, we predicted a developmental timepoint in which the MVA pathway regulates proliferation and YAP signaling in embryonic cardiomyocytes. This study exemplifies how our proteomic dataset can be used to identify and investigate pathways essential for embryonic heart development.

## DISCUSSION

Here we used high-throughput quantitative proteomics to measure temporal changes in the cardiac proteome at eight critical stages of embryonic heart development. We identified ∼7,300 proteins, assessed global temporal changes in protein expression, identified cardiac protein interaction networks, and linked protein dynamics with molecular pathways. In addition, we examined the temporal protein expression patterns of 185 CHD-associated proteins. Finally, we provide evidence of a function for the mevalonate (MVA) pathway in embryonic cardiomyocytes, a demonstration of how our dataset can be harnessed to investigate important molecular pathways that operate in heart development.

Delineating the temporal expression of protein networks and cellular pathways during critical stages of heart development is essential to understand the molecular mechanisms that drive cardiac morphogenesis. Our pairwise (E9.5 versus E11.5) and global temporal profiling analyses revealed unique biological pathways that are significantly upregulated early in cardiac development, including chromosome organization, cell-cycle, ribosome biogenesis, and septum development (Figure 3, 4). Intriguingly, within the chromosome organization pathway, we identified a network of proteins associated with the BAF, MOZ/MORF and Polycomb repressive chromatin-modifying complexes. Considerable research has shown that chromatin-modifying proteins are critical for heart development, and mutations in chromatin modifiers are major contributors to CHD (Jin et al., 2017). Further, the BAF, MOZ/MORF and Polycomb repressive complexes are known regulators of differentiation and maturation of the embryonic heart (He et al., 2012; Hota and Bruneau, 2016; Shirai et al., 2016; Vanyai et al., 2015; Voss et al., 2012).

Each of these multi-protein complexes contains a set of core and accessory protein subunits and the unique composition of each complex influences their molecular function. Interestingly, the BAF complex regulates cardiac differentiation decisions by systematically altering temporal changes in the subunit composition of the complex (Hota et al., 2019). Changes in subunit protein abundance constitute a primary mechanism that drives alterations in BAF complex composition. In agreement with these findings, we found that the BAF components SMARCD1, SMARCB1 and SMARCA1 were significantly upregulated at E9.5 versus E11.5, whereas other BAF subunits (SMARCA4, SMARCA2, SMARCD3) were not differentially expressed between these ages. These findings suggest that BAF complex subunits display distinct temporal expression profiles during embryonic heart development (Figure S4, TableS2). These findings highlight how our comprehensive proteomics dataset can be used to examine temporal protein expression dynamics of essential protein complexes during cardiac morphogenesis.

Aberrant ribosome biogenesis is associated with many congenital disorders referred to as ribosomopathies, some of which are present with congenital heart defects (Farley-Barnes et al., 2019; Venturi and Montanaro, 2020). Interestingly, our data revealed that a large number of proteins involved in ribosome biogenesis had peak protein abundance at early time points, then decreased rapidly as cardiac development proceeded (Figure 3B, 4B, S3). Ribosome biogenesis is a key means to control total protein synthesis and thereby regulate important functions such as cell growth, proliferation, and differentiation. Temporal regulation of ribosome biogenesis, hence protein translation, coordinates specific developmental events in many tissue types (Chau et al., 2018; Sondalle et al., 2016; Trainor and Merrill, 2014). Recent studies with embryonic stem cell derived cardiomyocytes suggest that downregulation of ribosome biogenesis and global protein synthesis are required for cardiomyocyte maturation (Pereira et al., 2019). Taken together, our data imply that strict regulation of ribosome biogenesis is essential for heart development. In addition, these data highlight our dataset as a critical resource for investigating the function of ribosome biogenesis during heart development and disease.

Notably, our global analysis showed that 51.7% (1,964/3,799) of differentially expressed proteins peaked early in gestation (Clusters 1, 2, 3) (Figure 4A). Additionally, most of the CHD-associated proteins were found within Clusters 1-3 (Figure 5). These findings imply that perturbations in protein abundance at early development stages (E9.5-E11.5) would be particularly detrimental to embryonic heart development. Further investigation of less studied proteins within these groups will provide an essential mechanistic understanding of cardiac development.

Proteins and cellular pathways enriched at later stages of embryonic heart development (E13.5-E16.5) reflect molecular mechanisms involved in the completion of multiple critical developmental events, including septation of the chambers, remodeling of the cardiac valves, and maturation of the myocardium. Our data showed overwhelming representation of proteins associated with metabolic pathways that are upregulated as the heart matures, including pathways of energy metabolism, fatty acid oxidation, and oxidative phosphorylation (Figure 3C, 3F, 4B, S5, S7). During embryogenesis, rapid growth and increased contractility of the heart are accompanied by drastic changes in cardiac metabolic activity. This metabolic change is best exemplified by the shift from glucose metabolism to oxidative phosphorylation that initiates during mid-to-late-gestation (Baker and Ebert, 2013; Gibb and Hill, 2018; Zhao et al., 2019). Interestingly, although we observed many metabolic proteins with peak abundances at late stages (i.e., E16.5) of heart development (Figure 4B), our data revealed a significant increase in proteins involved in glycolysis and fatty acid oxidation as early as E11.5 (Figure 3C). Alterations in cardiac metabolism are not only important for increased energy demands as the heart grows; they also influence critical developmental processes during heart morphogenesis. For example, alterations in glucose metabolism during embryogenesis result in several cardiac defects, such as impaired heart looping, hypoplasia of the myocardium, septal defects, and embryonic lethality (Gibb and Hill, 2018; Menendez-Montes et al., 2016; Smoak, 2002). Similarly, perturbations in oxidative phosphorylation are associated with severe cardiac defects and embryonic lethality (Kasahara et al., 2013; Zhao et al., 2019). Although temporal changes in cardiac metabolism are clearly essential for proper heart development, the molecular mechanisms of this process are understudied. Our dataset provides a foundation for examining the temporal dynamics of these pathways in heart development. Furthermore, alterations in cellular metabolism appear to drive cardiomyocyte proliferation during regeneration (Honkoop et al., 2019; Magadum et al., 2020). Therefore, our dataset will also be a valuable resource for the study of cardiac regeneration.

Investigation of proteins that are dynamically expressed during mid-gestation (Cluster 4, 5, and 6) revealed a function for the MVA pathway in heart development. The MVA pathway is an essential metabolic pathway composed of two branches that regulate cholesterol biosynthesis and protein prenylation. Cholesterol is important for cellular membrane structure, steroid hormone production, and signal transduction. Prenylation is a post-translational modification that facilitates the localization of proteins to membranes and regulates downstream signaling pathways. The MVA pathway has a well-established role in adult heart function and disease whereby alterations in the pathway are associated with many adult cardiovascular diseases including cardiac hypertrophy, fibrosis, and endothelial cell dysfunction (Table 1) (Yeganeh et al., 2014; Zhang et al., 2021). However, relatively little is known about the function of this pathway in cardiac development. Our work revealed new insights into the temporal expression dynamics of key proteins within the MVA pathway during heart development. Most of the core enzymes in the MVA pathway showed peak protein abundance between E11.5 and E13.5 (Figure 6A, 6B). Surprisingly, proteins specific to each downstream branch of the MVA pathway displayed divergent protein expression patterns. Proteins involved in cholesterol biosynthesis were upregulated at early stages of embryonic heart development (Cluster 2), whereas proteins that mediate prenylation were most abundant at later stages (Cluster 8) (Figure 6C, 6D and 6E). This finding suggested that the two branches of the MVA pathway are temporally controlled, and each branch contributes to different cardiac developmental processes.

Chen et al. reported that, in the embryonic heart, conditional loss of geranylgeranyl pyrophosphate synthase (GGPS1^KO^), an MVA pathway enzyme that regulates the production of isoprenoids used for protein prenylation, resulted in impaired ventricular maturation and embryonic lethality by E12.5 (Chen et al., 2018). Their report corroborates our findings that the MVA pathway core proteins and GGPS1 were highly expressed at this developmental time point (Figure 6A, 6D). Although a mechanistic explanation of the cardiac phenotype in GGPS1^KO^ mice is unclear, researchers suggested that impaired cardiomyocyte proliferation may be a contributor. Furthermore, a recent study with human pluripotent stem cell-derived cardiomyocytes identified the MVA pathway as a regulator of cardiomyocyte proliferation (Mills et al., 2019). Our studies using isolated E12.5 cardiomyocytes are the first to demonstrate that the MVA pathway directly regulates embryonic cardiomyocyte proliferation (Figure 7B, 7C).

Previous research, primarily in cancer studies, showed that the MVA pathway’s control of cell proliferation can be mediated by regulation of YAP/TAZ signaling (Gong et al., 2019; Mullen et al., 2016; Sorrentino et al., 2014; Wang et al., 2014). More recently, Chong et al. showed that MVA pathway regulation of vascular endothelial proliferation was mediated by control of YAP signaling (Chong et al., 2021). This finding is of particular significance because YAP/TAZ signaling regulates cardiomyocyte proliferation, embryonic heart morphogenesis, and cardiac regeneration (Chen et al., 2020; Moya and Halder, 2019). Our studies are the first to show that inhibition of the MVA pathway impairs YAP nuclear localization in embryonic cardiomyocytes. Therefore, we suggest that YAP regulation is a mechanism by which the MVA pathway controls embryonic cardiomyocyte proliferation.

Overall, we have generated the most comprehensive dataset of the murine embryonic heart proteome. Our temporal cardiac proteome profiling and functional studies of the MVA pathway exemplify how our quantitative proteomics dataset can be used to identify and investigate molecular pathways that are essential for embryonic heart development. This work provides a critical resource to study fundamental mechanisms of cardiac development and serves as a “blueprint” for investigating changes in the cardiac proteome that are associated with congenital heart disease.

## SUPPLEMENTAL FIGURE LEGENDS

**Figure 1 Supplemental:**
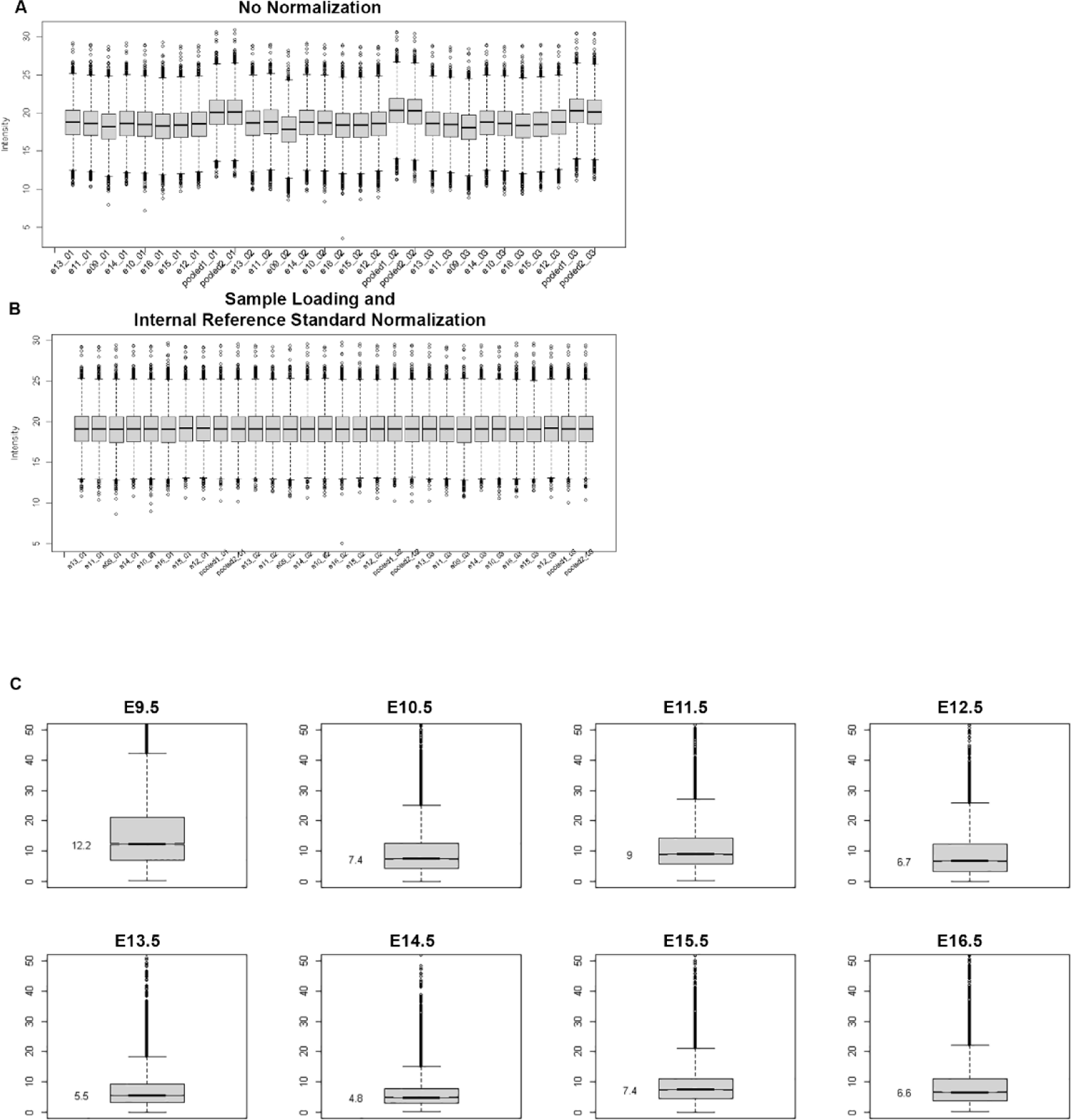
Protein abundance normalization and analysis of variance. (A) Mean ion intensities of each sample prior to normalization. (B) Mean ion intensities of each sample after sample loading and internal reference scaling normalization. (C) Protein abundance coefficient of variance across all samples.

**Figure 2 Supplemental:**
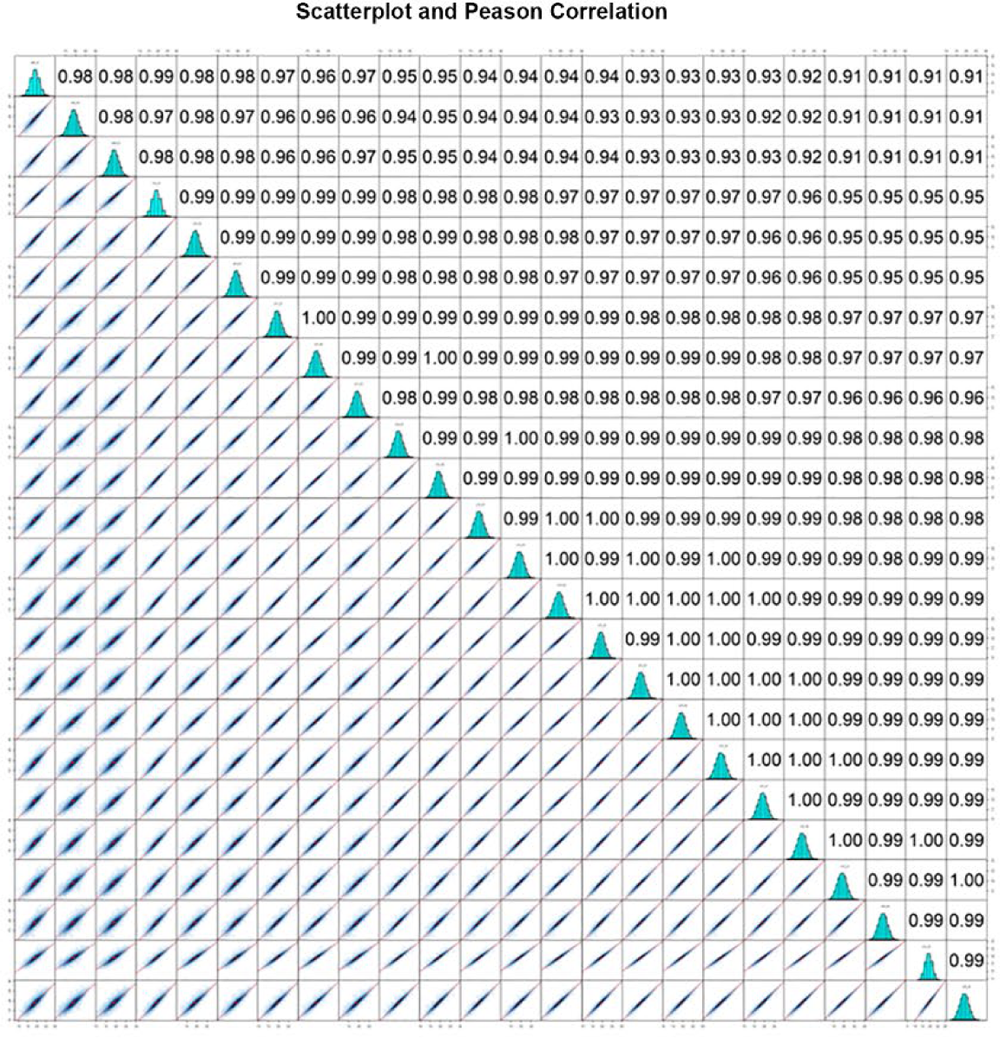
Scatter plots and Pearson correlation. Scatter matrices show pairwise Pearson correlation between all biological replicates for each age.

**Figure 3 Supplemental:**
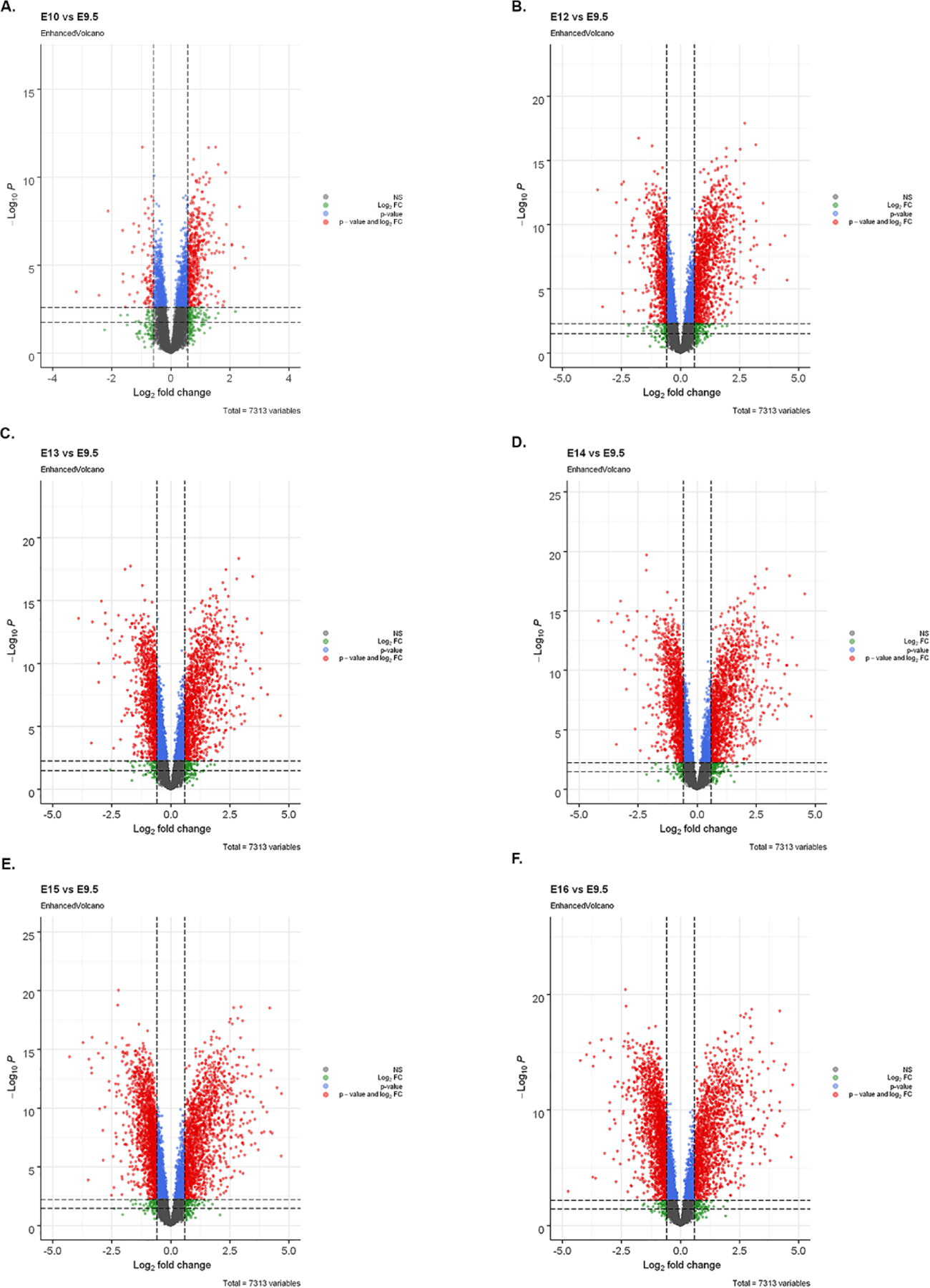
Differential proteins analysis at each age (versus E9.5). Related to Figure 1 and 4. Volcano plots showing proteins differentially expressed at E10.5 (A), E12.5 (B), E13.5 (C), E14.5 (D), E15.5 (E), E16.5 (F) compared to the E9.5 timepoint. Plots are shown as Log2 fold change (x-axis) plotted against the -log10 of the p-value (y-axis). Red dots represent proteins that were considered significantly different based on fold change (|Log_2_FC|≥0.585) and p-value (p-value ≤ 0.05) parameters. Blue dots represent proteins that only meet p-value parameters. Green dots represent proteins that only meet fold change parameters. Black dots represent proteins that are not differentially expressed.

**Figure 4 Supplemental:**
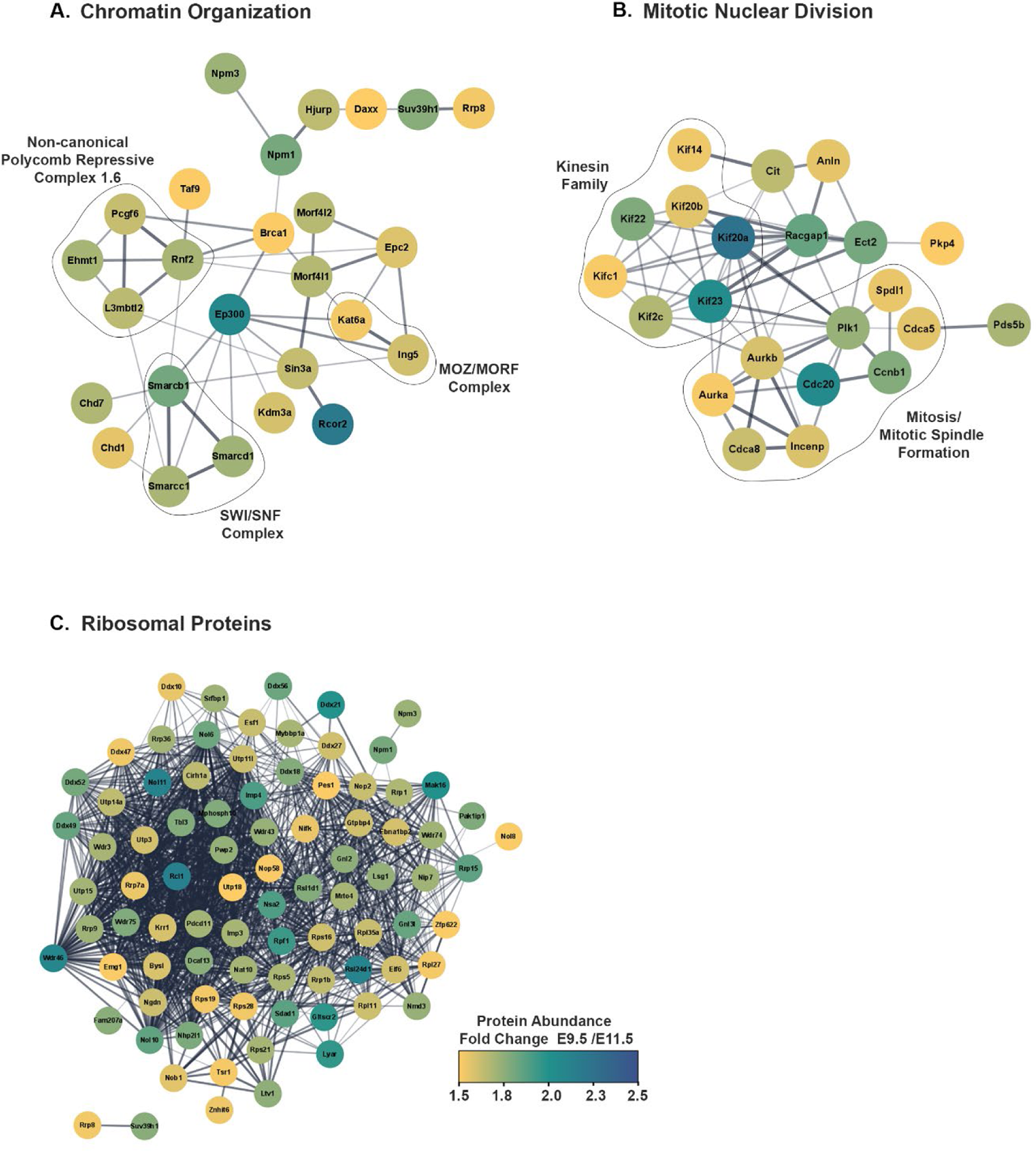
Proteins networks upregulated at E9.5 compared to E11.5. Related to Figure 3A and 3B. STRING network based on physical interactions of proteins upregulated at E9.5. Kappa score ≥ 0.4. Node color indicates fold change compared to E11.5.

**Figure 5 Supplemental:**
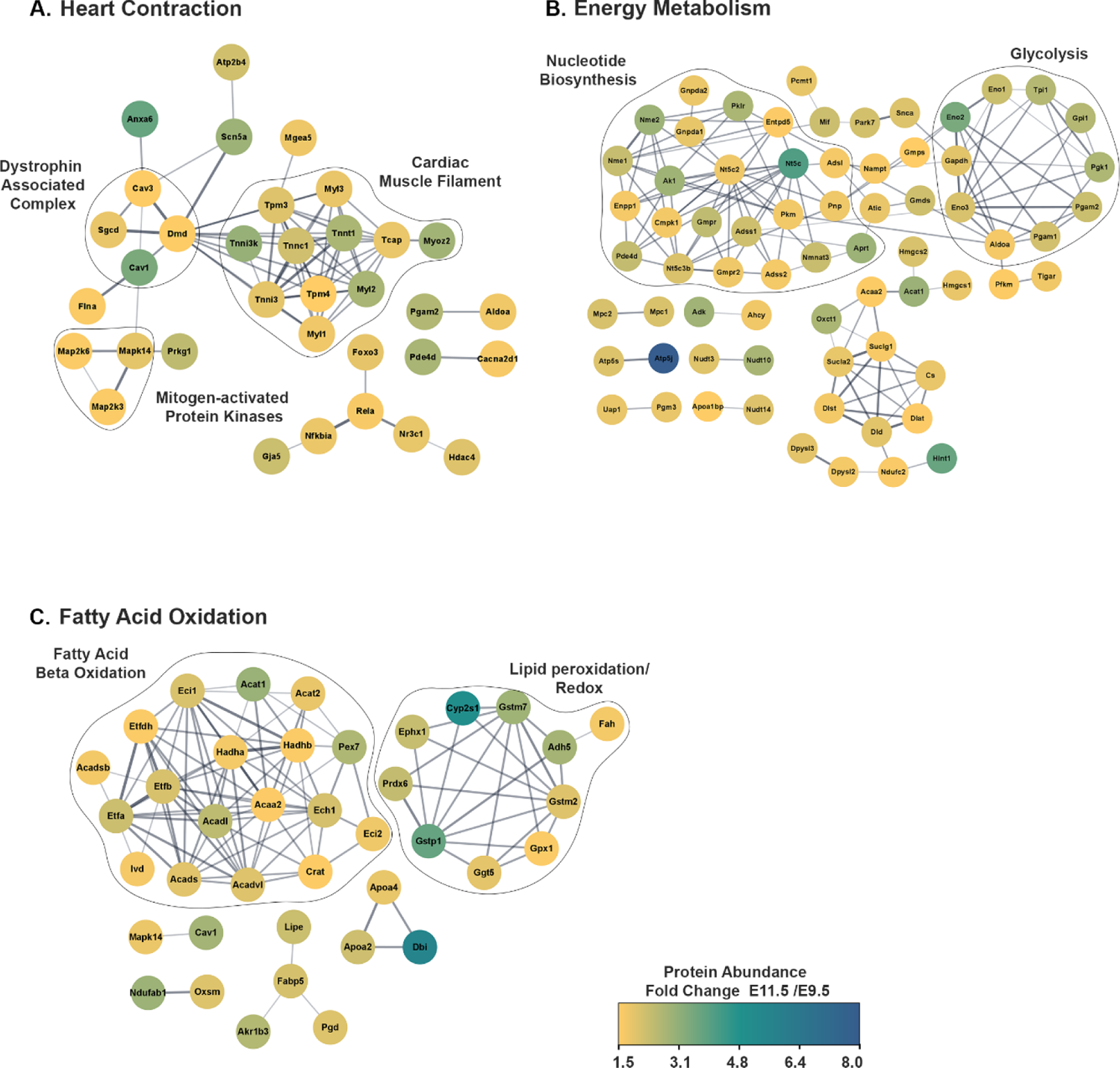
Proteins networks upregulated at E11.5 compared to E9.5. Related to Figure 3A and 3C. STRING network based on physical interactions of proteins upregulated at E11.5. Kappa score ≥ 0.4. Node color indicates fold change compared to E9.5.

**Figure 6 Supplemental:**
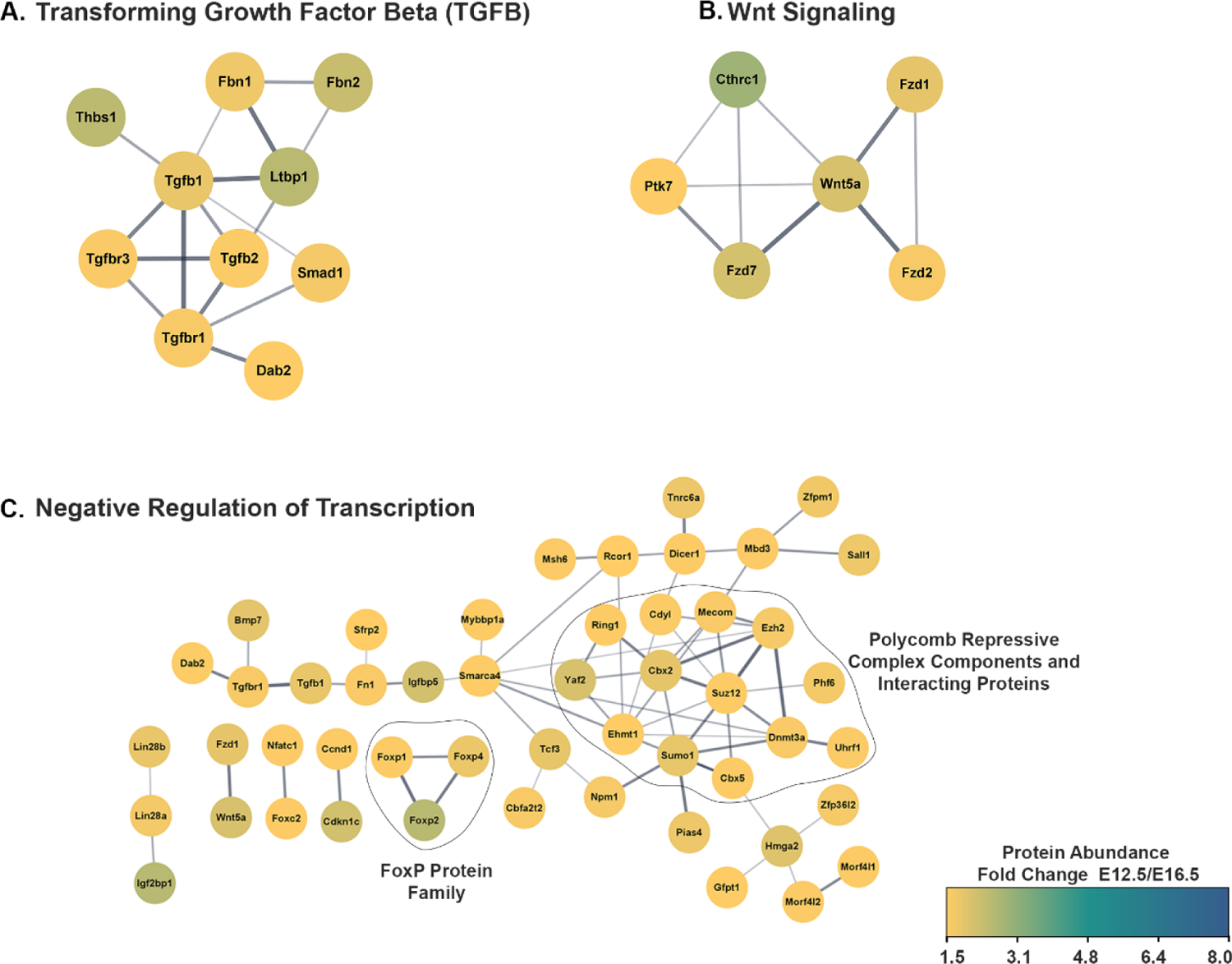
Proteins networks upregulated at E12.5 compared to E16.5. Related to Figure 3D and 3E. STRING network based on physical interactions of proteins upregulated at E12.5. Kappa score ≥ 0.4. Node color indicates fold change compared to E16.5.

**Figure 7 Supplemental:**
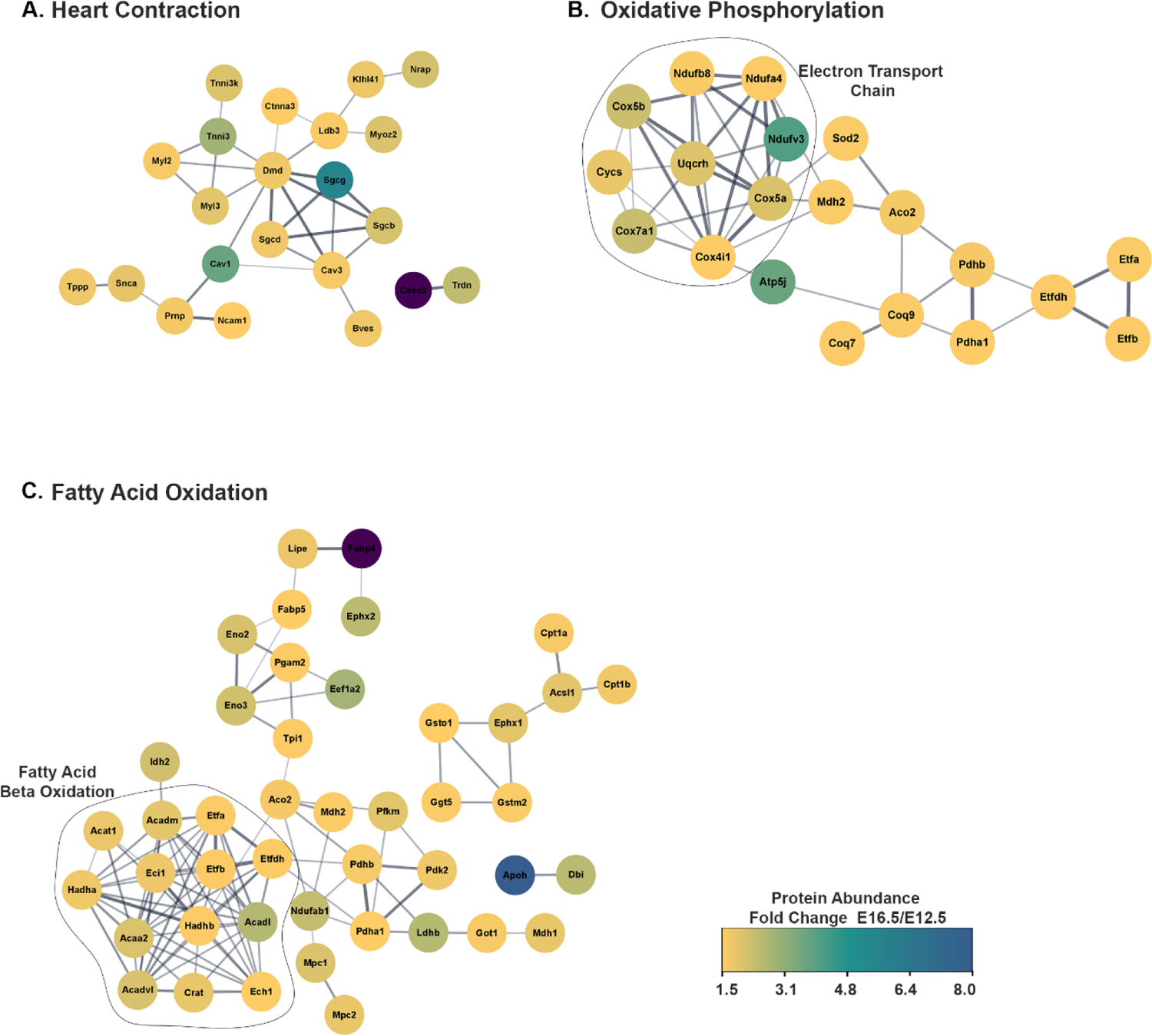
Proteins networks upregulated at E16.5 compared to E12.5. Related to Figure 3D and 3F. STRING network based on physical interactions of proteins upregulated at E16.5. Kappa score ≥ 0.4. Node color indicates fold change compared to E12.5.

**Supplementary Table 1:** Master table of all identified and quantified proteins.

**Supplementary Table 2:** Statistical analysis of differentially expressed proteins.

**Supplementary Table 3:** Hierarchical clustering heatmap input and computed values. Related to Figure 4A and 4B.

## METHODS

### Animal use

The Institutional Animal Care and Use Committee of the University of North Carolina approved all animal procedures and protocols, conformed to the Guide for the Care and Use of Laboratory Animals. C57/Bl6 male and female mice breed for timed matings were obtained from The Jackson Laboratory (Stock No: 000664).

### Sample preparation for proteomics analysis

Pregnant dams were sacrificed and subjected to transcardial perfusion with heparinized phosphate buffer saline to prevent blood coagulation. Embryonic hearts were dissected into HEPES buffer (20 mM HEPES, 1.2% PVP, pH 7.4) and snap frozen in liquid nitrogen, and stored at −80°C. To perform the protein extraction, hearts were thawed on ice and resuspended in lysis buffer (50 mM Tris-HCl pH 8.0, 100 mM NaCl, 0.5 mM EDTA, 2% SDS) supplemented with protease inhibitor and phosphatase inhibitors 2 and 3 (Sigma). Samples were subjected to dounce homogenization. Subsequently, samples were heated at 95°C for 5 minutes followed by sonication. Heating and sonication steps were repeated three times. Samples were centrifuged at 2000 x g for 5 minutes at room temperature to remove insoluble debris. Protein concentrations were determined by BCA assay. Fifty µg of protein was reduced and alkylated with 20 mM tris(2-carboxyethyl) phosphine and 20 mM chloroacetamide, respectively, for 20 minutes at 70°C. Samples were then precipitated using a methanol/chloroform protocol. Protein pellets were snap frozen in liquid nitrogen and stored at −80°C. Three biological replicates were collected for each embryonic age.

Protein pellets were reconstituted with 2 M urea in 50 mM ammonium bicarbonate, reduced with 5 mM DTT at 56°C for 30 minutes, then alkylated with 15 mM iodoacetamide at room temperature in the dark for 45 minutes. The samples were subjected to digestion with LysC (Wako) for 2 h and trypsin (Promega) overnight at 37°C at a 1:50 enzyme: protein ratio. The resulting peptide samples were acidified, desalted using Thermo desalting spin columns, then the eluates were dried via vacuum centrifugation. Peptide concentration was determined using Pierce Quantitative Colorimetric Peptide Assay. Fifteen µg of each sample was reconstituted with 50 mM HEPES pH 8.5, then individually labeled with 90 µg TMT 10plex reagent (Thermo Fisher) for 1 hour at room temperature. A pooled sample was created by combining a small amount of each sample then split into two aliquots, which were each labeled with two different TMT tags and used in all three TMT sets. Prior to quenching, the labeling efficiency was evaluated by LC-MS/MS analysis of a pooled sample consisting of 1 ul of each sample. After confirming >98% efficiency, samples were quenched with 50% hydroxylamine to a final concentration of 0.4%. Labeled peptide samples were combined, desalted using Thermo desalting spin column, and dried via vacuum centrifugation. The dried TMT-labeled samples (three TMT sets total) were fractionated using high pH reversed phase HPLC (Mertins et al., 2018). Briefly, the samples were offline fractionated over a 90 min run into 96 fractions by high pH reverse-phase HPLC (Agilent 1260) using an Agilent Zorbax 300 Extend-C18 column (3.5-µm, 4.6 × 250 mm) with mobile phase A containing 4.5 mM ammonium formate (pH 10) in 2% (vol/vol) LC-MS grade acetonitrile, and mobile phase B containing 4.5 mM ammonium formate (pH 10) in 90% (vol/vol) LC-MS grade acetonitrile. The 96 resulting fractions were concatenated in a noncontinuous manner into 24 fractions. The 24 fractions were dried via vacuum centrifugation.

### LC/MS/MS analysis

Three sets of 24 fractions were analyzed by LC/MS/MS using an Easy nLC 1200 coupled to an Orbitrap Fusion Lumos Tribrid mass spectrometer (Thermo Scientific) using a multi-notch MS3 method (McAlister et al., 2014) (reference: https://pubmed.ncbi.nlm.nih.gov/24927332/). Samples were injected onto an Easy Spray PepMap C18 column (75 μm id × 25 cm, 2 μm particle size) (Thermo Scientific) and separated over a 120 min method. The gradient for separation consisted of 5–42% mobile phase B at a 250 nl/min flow rate, where mobile phase A was 0.1% formic acid in water and mobile phase B consisted of 0.1% formic acid in 80% ACN. The Lumos was operated in SPS-MS3 mode with a cycle time of 3s. Resolution for the precursor scan (m/z 350–1600) was set to 120,000 with a AGC target set to standard and a maximum injection time of 50 ms. MS2 scans consisted of CID normalized collision energy (NCE) 30; AGC target set to standard; maximum injection time of 50 ms; isolation window of 0.7 Da. Following MS2 acquisition, MS3 spectra were collected in SPS mode (10 scans per outcome); HCD set to 65; resolution set to 50,000; scan range set to 100-500; AGC target set to 200% with a 105 ms maximum inject time.

### TMT-MS data processing

Raw data files were processed using Proteome Discoverer version 2.4, set to ‘reporter ion MS3’ with ‘10plex TMT’. Peak lists were searched against a reviewed Uniprot mouse database (downloaded May 2019 containing 17,457 sequences), appended with a common contaminants database, using Sequest HT within Proteome Discoverer. All fractions were searched with up to two missed trypsin cleavage sites, fixed modifications: TMT6plex peptide N-terminus and Lys, carbamidomethylation Cys, dynamic modification: N-terminal protein acetyl, oxidation Met. Precursor mass tolerance of 5ppm and fragment mass tolerance of 0.4 Da. Peptide false discovery rate was set to 1%. Reporter abundance based on intensity, SPS mass matches threshold set to 50, and razor and unique peptides wre used for quantitation. Data were further analyzed in R.

### Protein identification, quantification, and statistical analysis

Proteins selected for quantification were filtered based on the following criteria: 1) 1% False discovery rate 2) greater than two unique peptides detected, 3) at least one unique peptide is present in each of the six pooled reference channels and 4) the protein was detected in all three biological replicates in at least one sample group. Using the final list of filtered proteins (N = 7313) imputation was performed with the R function QRILC in the imputeLCMD package (https://rdrr.io/cran/imputeLCMD/).

Sample loading (SL) and internal reference scaling (IRS) normalization was performed as previously described (Plubell et al., 2017). Briefly, to correct for SL differences, reporter ion intensities were normalized within each 10-plex experiment by multiplying a global scaling factor to adjust the total ion intensity for each channel. To normalize reporter ion intensities between each TMT 10-plex (3 total) an IRS factor was calculated. For each protein, three reference intensities are calculated using the pooled reference samples from each TMT 10-plex. The three reference intensities are then averaged, and an IRS factor is computed for each protein. The IRS factor was then used to adjust reporter ion intensities for each protein in each TMT experiment.

To determine differential protein abundances across each age, pairwise statistical testing (versus E9.5) was performed using DEqMS (https://bioconductor.org/packages/release/bioc/html/DEqMS.html). Proteins with a |Log_2_FC| ≥ 0.585 (FC, fold change) and spectral count (sca) adjusted p-value ≤ 0.05 at a least one age were considered differentially expressed. Functional over-representation of biological processes analysis of differentially expressed proteins was performed with using clusterProfiler package (https://bioconductor.org/packages/release/bioc/html/clusterProfiler.html). The principal component analysis (PCA) and heatmap generation were performed on the web-based version of ClustVis (https://biit.cs.ut.ee/clustvis_large/). Volcano plots were constructed using EnhancedVolcano (https://bioconductor.org/packages/release/bioc/html/EnhancedVolcano.html)

Protein interaction network diagrams incorporating known functional and physical protein interactions were generated using Cytoscape with the STRING database plugin (Kappa score ≥ 0.4). For functional analysis of proteins identified as differentially expressed from pairwise comparisons of E9.5 vs E11.5 and E12.5 vs E16.5, the Cluego plugin in Cytoscape was used to generate pie charts of significantly enriched (p-value ≤ 0.001) gene ontology terms. Pie charts represent the percent of gene ontology terms per group.

### Cardiomyocyte isolation and treatment

Cardiomyocytes were isolated from E12.5 embryonic hearts according to a previously published protocol (Ehler et al., 2013). Briefly, hearts were harvested and washed in 1X PBS supplemented with 20 mM BDM. Hearts were then minced and incubated in isolation medium (20 mM BDM, 0.0125% trypsin in HBSS (without Ca2+, Mg2+) overnight at 4°C. The following day the predigested hearts were transferred to digestion medium (20 mM BDM, 1.5 mg/ml Roche Collagenase/Dispase enzyme mix in L15 medium) and were incubated at 37°C for 20 minutes with gentle shaking. Tissue fragments were triturated 20 times and the cell suspension was strained. Cells were centrifugated at 800 x g for 5 minutes. The supernatant was removed, and cells were resuspended in 10 ml of plating medium (65% DMEM high glucose, 19% M-199, 10% horse serum, 5% fetal calf serum, 1% penicillin/streptomycin) and plated into a 10 cm cell culture dish. Cells were incubated at 37°C for 3 hours to allow adherence of non-cardiomyocyte cells. After incubation, the media containing the non-adherent cardiomyocytes was removed and transferred to a 15 ml conical Falcon tube. Cells were centrifuged at 800 x g for 10 minutes and cells were resuspended in 2 ml of plating media. Cells were counted and approximately 7.5 x 10^4^ cells were plated into collagen coated, 8-well chamber slides. The following day the cells were change into maintenance media (78% DMEM high glucose, 17% M-199, 4% horse serum, 1% penicillin/streptomycin). The following day cells were treated with maintenance medium containing 10 µM Simvastatin for 24 hours.

### Immunohistochemistry

For immunochemical analysis of isolated cardiomyocytes, cells were fixed with 1% paraformaldehyde in 1X PBS for 15 minutes at room temperature, permeabilized with 0.2 % Triton X-100 in 1X PBS for 10 minutes at room temperature and incubated with blocking buffer (3% Normal Goat serum, 0.05% Triton X-100 in 1X PBS) for 1 hour at room temperature. Cells were then incubated with primary antibody in antibody buffer (1% Normal Goat serum, 0.05% Triton X-100 in 1X PBS) overnight at 4°C. The following day cells were incubated in secondary antibody diluted in IHC antibody buffer for 1 hour at room temperature, and subsequently counterstained with 200 ng/ml 4′,6-diamidino-2-phenylindole (DAPI) for 10 minutes at room temperature.

### Image quantitation

For quantitation of the percent of Ki67-positive cardiomyocytes and YAP nuclear localization, ten, 20x images were analyzed per well. At least, three biological replicates were performed for each experiment and at least 300 cells were analyzed for each replicate. CellProfiler was used for analysis and cell counts. Data were compared for statistical significance by Student’s *t*-test.

| REAGENT or RESOURCE | SOURCE | IDENTIFIER |
| --- | --- | --- |
| **Antibodies** |  |  |
| Tropomyosin (CH1) | DSHB | CH1 |
| Ki67 | abcam | ab15580 |
| YAP | Cell Signaling | 49125 |
| Alexa Flour 488 Goat anti-Mouse | Invitrogen | A11001 |
| Alexa Flour 546 Goat anti-Rabbit | Invitrogen | A11035 |
| **Biological samples** |  |  |
| **Chemicals, peptides, and recombinant proteins** |  |  |
| TCEP | Thermo | 77720 |
| Chloroacetamide | Sigma | 22790 |
| Protease inhibitor | Sigma | P8340 |
| Phosphatase inhibitor cocktail 2 | Sigma | P5726 |
| Phosphatase inhibitor cocktail 3 | Sigma | P0044 |
| BDM (2,3-Butanedione monoxime) | Sigma | B0753 |
| Collagenase/Dispase | Sigma | 1026963800 |
| 1x trypsin solution (0.25%) with EDTA | Gibco | 25300-054 |
| DMEM high glucose | Sigma | D6429 |
| M-199 | Fisher | MT10060CV |
| Fetal bovine serum (FBS) | VWR | 97068-85 |
| Horse serum | Gibco | 16050-122 |
| Leibovitz L-155 | Corning | 10-045-CV |
| 0.1% collagen solution | Sigma | C8919 |
| Normal goat serum | Sigma | 526 |
| **Critical commercial assays** |  |  |
| TSA Cyanine 3 Kit | Akoya Biosciences | NEL704A001KT |
| BCA Assay | Thermo | 23225 |
| **Deposited data** |  |  |
| MS/MS Raw files and MaxQuant analysis files | This paper | ProteomeXchange: |
| **Experimental models: Organisms/strains** |  |  |
| Mouse strain wild type C57BL/6 | The Jackson Laboratory |  |
| **Software and algorithms** |  |  |
| CellProfiler | Broad Institute | https://cellprofiler.org/ |
| Biorender |  |  |
| Graphpad Prism |  |  |
| Proteome Discover 2.4 | Thermo Scientific | N/A |
| Cytoscape with ClueGO and STRING plugin |  |  |
| **Other** |  |  |

